# Systematic analysis of the R2R3-MYB family of transcription factors in *Camellia sinensis*: evidence for species-specific catechin biosynthesis regulation

**DOI:** 10.1101/2021.07.21.453189

**Authors:** Jingyi Li, Shaoqun Liu, Peifen Chen, Jiarong Cai, Song Tang, Wei Yang, Fanrong Cao, Peng Zheng, Binmei Sun

## Abstract

The R2R3-MYB transcription factor family regulates metabolism of phenylpropanoids in various plant lineages. Species-expanded or specific MYB transcription factors may regulate species-specific metabolite biosynthesis including phenylpropanoid-derived bioactive products. *C. sinensis* produces an abundance of specialized metabolites, which makes it an excellent model for digging into the genetic regulation of plant-specific metabolite biosynthesis. The most abundant health-promoting metabolites in tea are galloylated catechins, and the most bioactive of the galloylated catechins, epigallocatechin gallate (EGCG), is exclusively found in *C. sinensis*. However, the transcriptional regulation of galloylated catechin biosynthesis remains elusive. This study mined the R2R3-MYB transcription factors associated with galloylated catechin biosynthesis in *C. sinensis*. A total of 118 R2R3-MYB proteins, classified into 38 subgroups, were identified. R2R3-MYB subgroups specific to or expanded in *C. sinensis* were hypothesized to be essential to evolutionary diversification of tea-specific metabolites. Notably, nine of these *R2R3-MYB* genes were expressed preferentially in apical buds and young leaves, exactly where galloylated catechins accumulate. Three putative *R2R3-MYB* genes displayed strong correlation with key galloylated catechin biosynthesis genes, suggesting a role in regulating biosynthesis of epicatechin gallate (ECG) and EGCG. Overall, this study paves the way to reveal the transcriptional regulation of galloylated catechins in *C. sinensis*.

## Introduction

Tea from *Camellia sinensis*, along with coffee and cocoa, is one of the world’s three major non-alcoholic beverages. Worldwide, approximately two billion cups of tea are consumed daily (Drew et al., 2019; Yamashita et al., 2020). Tea production has amplified at an average annual rate of 3.35% in the last five years; by 2019, worldwide tea production reached 6.49 million tons on 5.07 million hectares (Food and Agriculture Organization of the United Nations statistics, https://www.fao.org/faostat/). Tea was used first as a food in ancient China, then it served as a medicine to prevent and cure common diseases before developing into a popular beverage (Abbas et al., 2017; Mondal et al., 2004). Nowadays, tea exhibits great commercial potential and has become a vital industry due to its health promoting properties and attractive distinct flavors (Chen et al., 2009).

The tea plant (*C. sinensis*) is rich in characteristic metabolites, such as polyphenols, amino acids, caffeine, and terpenes, that significantly contribute to its pleasant flavors and industrial and medical value (Wei et al., 2018). Catechins are the principal health-promoting bioactive compounds of tea. Catechins constitute 12 to 24% of the dry weight of young leaves, and account for more than 70% of the total polyphenols (Yang et al., 2012). Catechins in tea consist of a mixture of catechin (C), epicatechin (EC), gallatechin (GC), epigallocatechin (EGC), catechin gallate (CG), epicatechin gallate (ECG), gallatechin gallate (GCG) and epigallocatechin gallate (EGCG) (Asakawa et al., 2013). Among them, galloylated catechins ECG and EGCG are abundant in tea plants and account for more than 80% of total catechins (Kim et al., 2004; Liu et al., 2012). EGCG, exclusively present in *C. sinensis*, is the major bioactive component conferring the many health benefits of tea - it is anti-carcinogenic (Ahmad et al., 2000), anti-oxidative (Heim et al., 2002), anti-bacterial and anti-inflammatory (Taguri et al., 2004) and it prevents cardiovascular and cerebrovascular diseases (Yu et al., 2020). In addition, EGCG is widely used in food production on account of its strong antioxidative capacity(Nikoo et al., 2018).

To date, based on biochemical, physiological and genetic research, the biosynthesis pathway of catechins has become clear (Wei et al., 2018; Yang et al., 2012; Yu et al., 2021). Catechins are derived from the phenylpropanoid pathway and principally accumulate in apical shoots and young leaves. Galloylated catechin content is mainly regulated at the transcriptional level by catechin biosynthesis genes *dihydroflavonol reductase* (*DFR*), *anthocyanidin reductase* (*ANR*), *leucoanthocyanidin reductase* (*LAR*) and *serine carboxypeptidase-like acyltransferases* (*SCPLs)* (Ashihara et al., 2010; Eungwanichayapant et al., 2009; Punyasiri et al., 2004; Singh et al., 2008; Wei et al., 2018). However, few studies have focused on the network(s) that regulate catechin biosynthesis, especially galloylated catechins.

R2R3-MYB transcription factors (TFs) comprise the largest family of TFs in advanced plants (Ambawat et al., 2013). In addition to possessing two imperfect MYB repeats (R2 and R3), the R2R3-MYB TFs maintain a highly conserved N-terminal MYB DNA-binding domain and an activated or repressed C-terminal domain (Dubos et al., 2010; Jiang et al., 2004; Jin and Martin, 1999; Kranz et al., 1998; Lipsick, 1996; Martin et al., 1997). The R2R3-MYB family is widely involved in plant growth and development, primary and secondary metabolism, hormone signal transduction, cellular proliferation and apoptosis, as well as disease and abiotic stress response (Li et al., 2017; Martin et al., 1997). Notably, the R2R3-MYB family plays an important role in positively or negatively regulating the biosynthesis of specialized metabolites, such as flavonoids (Hichri et al., 2011; Mehrtens et al., 2005), anthocyanin (Li et al., 2020; Li et al., 2017; Yu et al., 2020) and lignin (Bedon et al., 2007; Goicoechea et al., 2005).

Studies have found that new R2R3-MYB TFs emerged through species-specific duplication events (Soler et al., 2015). Species-specific evolved or expanded R2R3-MYB membership seems to confer functional diversification to organisms (Zhang et al., 2000). For example, the ancestral R2R3-MYB anthocyanin master regulator expanded into several homologous clusters within the grape (*Vitis* spp.) and maize (*Zea mays*) genomes, and differential expression of duplicated genes resulted in control of anthocyanin biosynthesis in different tissues (Jiu et al., 2021; Zhang et al., 2000). Some species-specific and expanded *R2R3-MYB* TFs govern specialized metabolite biosynthesis within lineages (Zhu et al., 2019). In *Capsicum*, five Solanaceae-specific MYB TF tandem genes duplicated in the *Cap1/Pun3* locus. *Capsicum* species have evolved placenta-specific expression of MYB31, which directly activates expression of capsaicinoid biosynthetic genes and results in production of genus-specialized metabolites. In *C. sinensis*, only a few R2R3-MYB transcription factors have a demonstrated role in regulation of phenylpropanoid biosynthesis. Specifically, *CsAN1*, *CsMYB6A* and *CsMYB75* regulate anthocyanin pigments in *C. sinensis* leaf (He et al., 2018; B. Sun et al., 2016; Wei et al., 2019). However, the *C. sinensis* specific and expanded R2R3-MYB TFs that are potential candidate regulators of galloylated catechins biosynthesis have still not been identified.

In this study, through performing a genome-wide analysis of the R2R3-MYB superfamily in *C. sinensis*, we compared the phylogenetic relationships between *C. sinensis* and other plant lineages. The gene structure, conserved motifs and transcript patterns were analyzed. Because of the importance of galloylated catechins in *C. sinensis*, we focused on the discovery of *R2R3-MYB* genes potentially involved in the regulation of biosynthesis of ECG and EGCG, especially EGCG which is unique to *C. sinensis*.

## Materials and Methods

### Plant materials

The ‘Lingtoudancong’ variety of *C. sinensis* was grown at South Agricultural University in Guangzhou, China. Apical buds, first leaves, second leaves, mature leaves, old leaves, stems and roots of ‘Lingtoudancong’ were sampled in spring of 2021. The samples of different tissues were immediately frozen in liquid nitrogen and stored at -80°C.

### Phylogenetic analysis

*C. sinensis* MYB protein sequences were retrieved from the Tea Plant Information Archive database (http://tpia.teaplant.org/). In total, 222 MYBs and MYB-related genes were predicted in the ‘Shuchazao’ genome, but only 118 of these had two consecutive and conserved repeats of the MYB domain. The R2R3-MYB protein sequences of *Arabidopsis thaliana* were obtained from the Arabidopsis Information Resource Archive database (https://www.arabidopsis.org/). The number of R2R3-MYB gene models identified by our methodology in the genome of *A. thaliana* (126) was the same as described in the literature (Dubos et al., 2010). The homologous genes of kiwifruit, coffee, cacao and grape were retrieved from the Plant Transcription Factor Database (http://planttfdb.gao-lab.org/) by performing a reverse BLAST search. All the R2R3-MYB sequences were aligned using ClustalX, and a neighbor-joining phylogenetic tree was constructed with 1,000 bootstrap replicates utilizing MEGAX (Kumar et al., 2016). A pairwise deletion method was chosen to dispose of the positions containing gaps or missing data in the sequences, and the delay divergent cutoff value was set to 30.

### Conserved motif analysis of R2R3-MYB

Functional motifs and conserved domains were identified with The MEME Suite tool (https://meme-suite.org/meme/) using the following parameters: site distribution, zero-or-one-site-per-sequence (ZOOPS) model; maximum number of motifs: 20; minimum motif width: 6; maximum motif width: 50; minimum number of sites per motif: 2; and maximum number of sites per motif: 118 (Bailey et al., 2009; Chen et al., 2021). The sequence logos of R2 and R3 repeats of the R2R3-MYB proteins were based on multiple sequence alignments and were visualized with WebLogo Version 2.8.2 (http://weblogo.berkeley.edu/logo.cgi). All obtained motifs were constructed and visualized using the Gene Structure View (Advanced) of the TBtools software (Chen et al., 2020).

### RNA-seq expression analysis

The RNA-seq data were downloaded from TPIA for transcript abundance analyses. The expression levels of the candidate *R2R3-MYB* TFs and catechin biosynthesis genes from different tissues of *C. sinensis* were used to generate a heatmap with TBtools software using the normalized method.

### RNA extraction and quantitative real-time PCR (qRT-PCR)

Total RNA of different tissues of ‘Lingtoudancong’ was extracted utilizing a Magen HiPure Plant RNA Mini kit B (R4151, Magen, China) according the manufacturer’s instructions. First-strand cDNA was synthesized using a HiScript III RT 1^st^ Strand cDNA Synthesis kit (R323-01, Vazyme, China) in a reaction volume of 20 μL. qRT-PCR was performed in a Bio-Rad CFX384 Touch^TM^ system. Each 10 μL reaction mixture was comprised of 4.4 μL qPCR SYBR Green Master Mix (Yeasen, China), 4.4 μL double distilled water, 0.2 μL of each primer (10 μmol/μL) and 1 μL of cDNA template. The reaction program was as follows: 95 °C for 5 min; then 39 cycles at 95 °C for 5 s and 60 °C for 30 s. A melting-curve analysis was carried out at 95 °C for 5 s, which was followed by a temperature increase from 60 °C to 95 °C. *Actin* (*TEA019484.1)* was used as the housekeeping gene. The relative expression of each gene was calculated with the 2^−ΔΔCt^ method (Livak and Schmittgen, 2001). The qRT-PCR primers were designed with the qPrimerDB-qPCR Primer Database (https://biodb.swu.edu.cn/qprimerdb/). Sequences of the primers are listed in Supplementary Table S2. Values were the means ± SDs of 3 replicates.

### Quantification of catechin contents

Reference standards of catechin (C), epicatechin (EC), epigallocatechin (EGC), epicatechin gallate (ECG), and epigallocatechin gallate (EGCG) were purchased from Shanghai Yuanye Bio-Technology Co., Ltd. (Shanghai, China). Apical buds, first leaves, second leaves, mature leaves, old leaves, stems and roots of ‘Lingtoudancong’ were ground into fine powders and freeze-dried. Approximately 0.2 g of each sample powder was extracted with 8 mL of 70% methyl alcohol (diluted with ultrapure water). After ultrasonic extraction for 30 min, the supernatant was collected by centrifugation. 1 mL of the liquid supernatant was filtered through a 0.22 μm Millipore membrane. The extracts were injected into an XSelect HSS C18 SB column (4.6 × 250 mm, 5 μm, Waters Technologies, USA). The catechin monomers were separated using 0.1% aqueous formic acid (A) and 100% acetonitrile (B) as mobile phases on a Waters Alliance Series HPLC system (Waters Technologies, USA). Detection was performed at 280 nm. Data were presented as the mean ± SD (n = 3).

### Correlation analysis of gene expression and metabolite accumulation

The correlation analysis among transcription factors, catechin biosynthesis genes and catechin monomer contents was performed via Pearson’s correlation coefficients. The R software was adopted to visualize the relationship directly. A correlation coefficient of >0.5 was considered to be a positively associated pair, and R<-0.5 was thought of as a negative correlation. In the diagram, blue represents a positive correlation, and red represents a negative correlation.

## Results

### Comparative phylogenetic analysis of the R2R3-MYB families in *C. sinensis* and *A. thaliana*

A total of 118 *R2R3-MYB* genes were identified in the *C. sinensis* genome after manual curation and exclusion of alternative transcripts. All identified *R2R3-MYB* genes from *C. sinensis* (118) were aligned with those of *A. thaliana* (126), and their evolutionary history was inferred by constructing a neighbor-joining phylogenetic tree (Fig. 1). The 118 *R2R3-MYB* genes of *C. sinensis* were named in light of the systematic naming rules of *A. thaliana*, except for *CsMYB1*, *CsMYB4a* and *CsAN1*, which had been functionally characterized previously (Li et al., 2017; Sun et al., 2016a; Yang et al., 2012) (Supplementary Table S1). In addition, as it is believed that genes which clustered together were considered to be in the same subgroup, and *A. thaliana* is, by far, the species for which the *R2R3-MYB* genes have been most extensively investigated, the 38 subgroups were classified by taking into account the topology of the tree and the bootstrap values (Fig. 1) and were named according to the classification of *A. thaliana* (Dubos et al., 2010). For new subgroups not previously proposed in *A. thaliana*, the subgroup was named after the known functionally characterized *A. thaliana* member.

**Figure 1.**
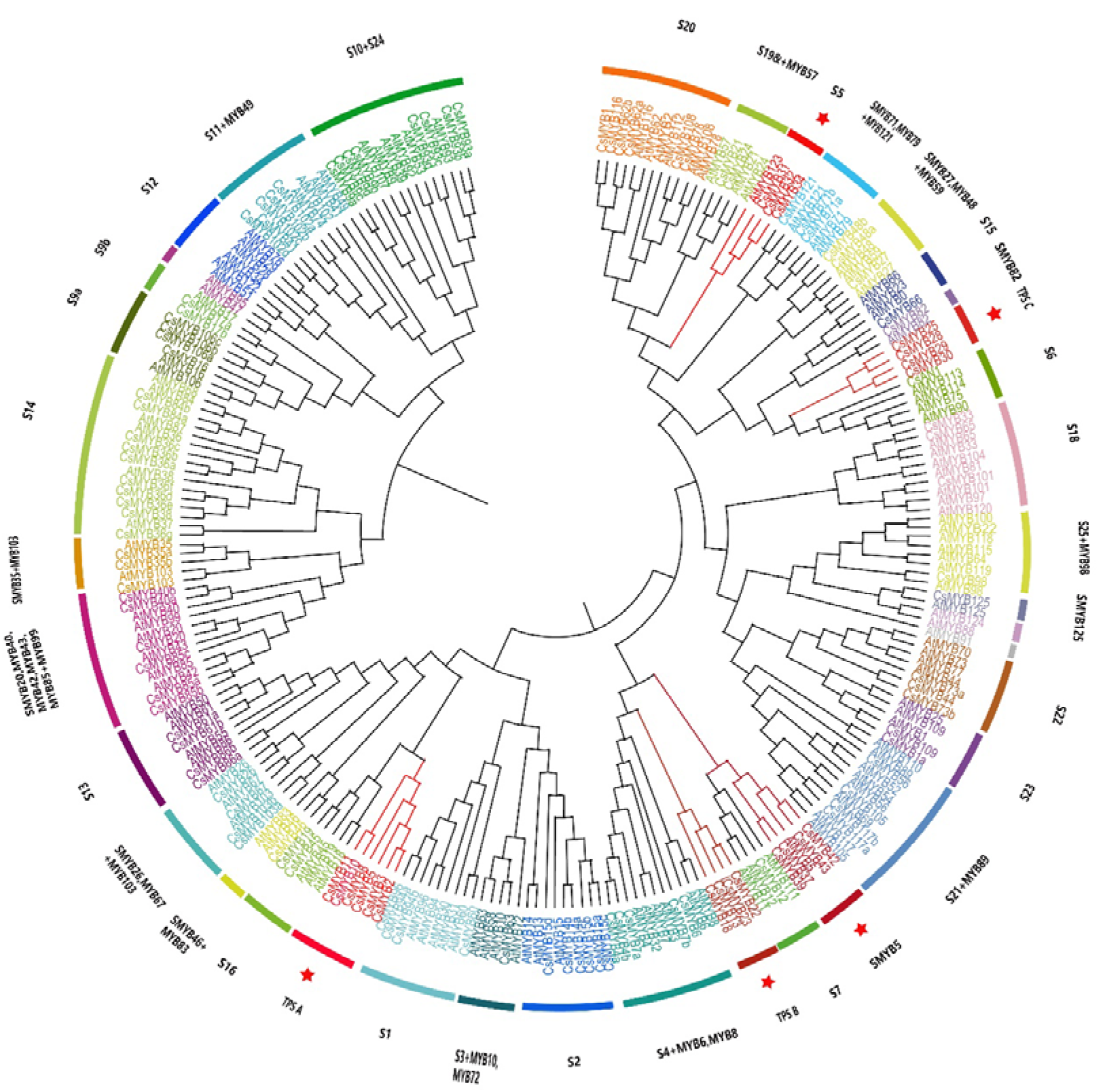
Phylogenetic analysis of the R2R3-MYB families in *C. sinensis* and *A. thaliana*. A neighbor-joining phylogenetic tree was constructed from 244 protein sequences including all R2R3-MYB proteins from *C. sinensis* (118) and *A. thaliana* (126). Subgroups within each clade were given a different color; meanwhile, the same color indicates the genes are in the same subgroup. Subgroup short names are included next to each clade to simplify nomenclature. Subgroups that evolved and expanded exclusively in the *C. sinensis* genome are highlighted in red and marked with a red star.

The majority of subgroups contained members from both species. However, S12 was an *Arabidopsis*-specific subgroup, containing only *R2R3-MYB* members from *A. thaliana*. The members of subgroup S12, *AtMYB28*, *AtMYB29*, *AtMYB76*, *AtMYB34* and *AtMYB5*1 regulate glucosinolate biosynthesis, a metabolite exclusive to the *Brassicaceae* family (Gigolashvili et al., 2007; Matus et al., 2008). In contrast, three subgroups contained *R2R3-MYB* TFs that were present only in *C. sinensis* without any homologues in *A. thaliana*. Therefore, their names we designated as Tea Preferential Subgroup A (TPSA), Tea Preferential Subgroup B (TPSB) and Tea Preferential Subgroup C (TPSC). Remarkably, SMYB5 and S5 subgroups were comprised of more tea plant *R2R3-MYB* members than *A. thaliana* members. For example, subgroup S5 had three *C. sinensis* members (*CsMYB31*, *CsMYB32* and *CsMYB34*), while *A. thaliana* contributed only one member, *AtMYB123/TT2.* Similarly, the SMYB5 subgroup had four members from *C. sinensis* (*CsMYB37*, *CsMYB39*, *CsMYB42* and *CsMYB43*), but just one member (*AtMYB5*) from *A. thaliana.* In *Arabidopsis*, the members of SMYB5 and S5 subgroups are involved in the phenylpropanoid pathway. *AtMYB123* controls the biosynthesis of proanthocyanidins (PAs) in the seed coat and *AtMYB5* was partially redundant with *AtMYB123* (Gonzalez et al., 2009). The expansion of SMYB5 and S5 subgroups in *C. sinensis* suggests that diversified regulation of polyphenols emerged during the speciation of *C. sinensis*.

Overall, the phylogenetic analysis results highlighted five special subgroups TPSA, TPSB, TPSC, S5 and SMYB5, which expanded in or were exclusively present in *C. sinensis*. Among these subgroups, a total of 21 *R2R3-MYB* TFs (TPSA (6), TPSB (4), TPSC (4), S5 (3) and SMYB5 (4)) attracted our attention and were selected for further analysis as potential candidates involved metabolic processes specific to *C. sinensis*.

### Functions inferred through phylogenetic analysis of the candidate subgroups in six plant species

To evaluate the 21 *R2R3-MYB* genes of the five tea-specific and tea-expanded subgroups TPSA, TPSB, TPSC, S5 and SMYB5, the homologous *R2R3-MYB* genes from kiwifruit, coffee, cacao, and grape were used to construct phylogenetic tree (Fig. 2). The number of *R2R3-MYB* TFs presented a trend of expansion in *C. sinensis* (21) similar to coffee (20), but greater than cocoa (11), kiwifruit (11) and grape (18). We surmised that the *C. sinensis* expanded *R2R3-MYB* function may have occurred through divergent evolution during speciation. For example, subgroup TPSA was expanded in tea (6) relative to coffee (3), cocoa (5), kiwifruit (5) and grape (1). In the TPSB subgroup, coffee (4), grape (4) and kiwifruit (3) had comparable numbers of members with *C. sinensis* (4), but only two homologous genes were found in cocoa. Subgroup TPSC contained homologous genes in all species except for kiwifruit. Remarkably, no isogenous genes of TPSA, TPSB or TPSC subgroups were present in *A. thaliana*, indicating that these *R2R3-MYB*s evolved only in tea plant and probably have novel functions.

**Figure 2.**
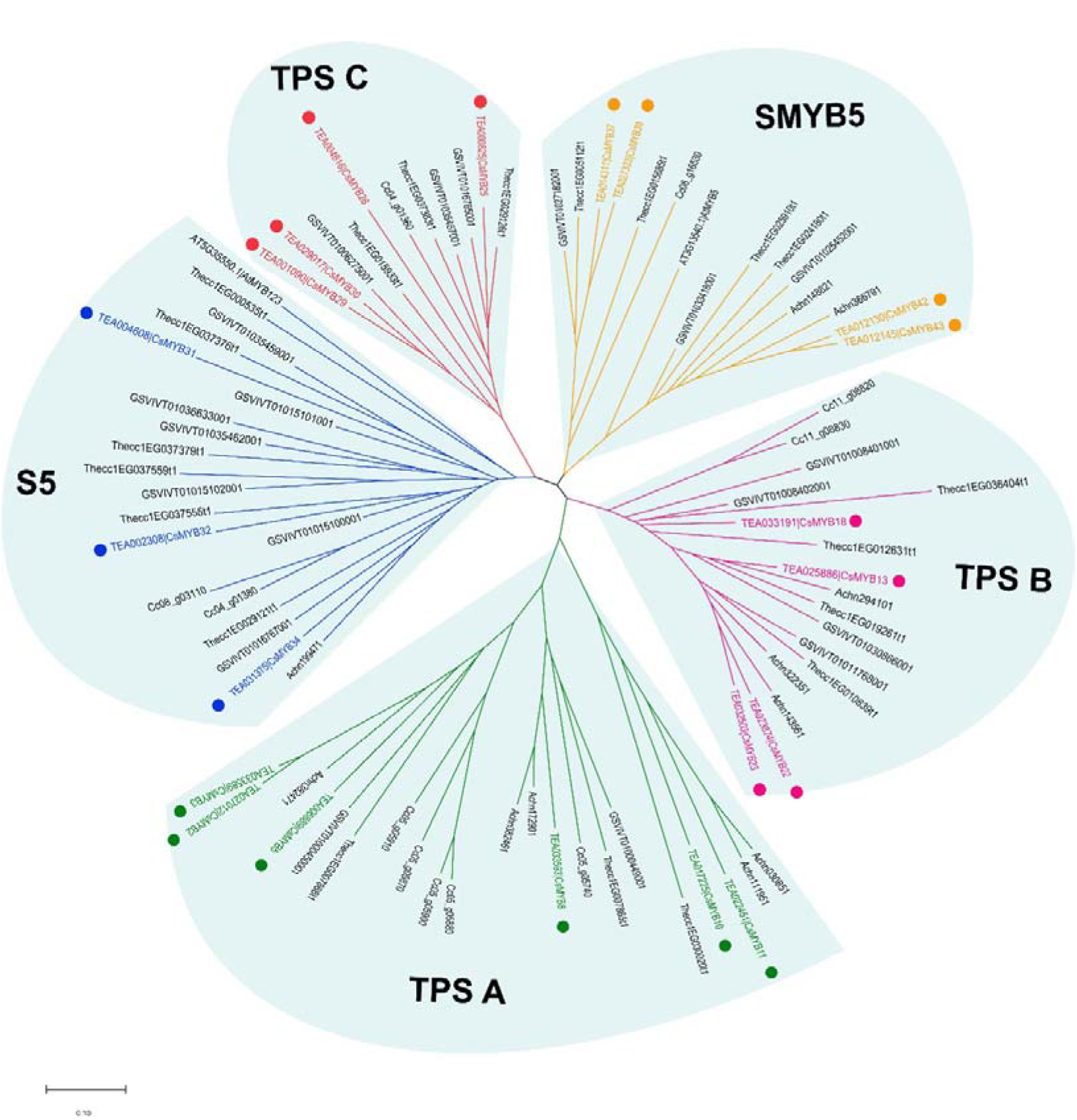
Phylogenetic analysis of the candidate subgroups in six plant species. A neighbor-joining phylogenetic tree was constructed with the R2R3-MYB proteins from *C. sinensis*, *Actinidia chinensis*, *Vitis vinifera*, *Theobroma cacao*, *Coffea canephora* and *Arabidopsis* genomes. Subgroup short names are indicated beside each clade.

To uncover the roles these *R2R3-MYB* genes serve, we searched for the functional characteristics of the selected *R2R3-MYB* genes from four close relative species. Only a few homologous genes (*GSVIVT01026868001*, *Achn38246* and *Achn172901* from the TPSA subgroup; *Achn143561*, *Achn322351* and *GSVIVT0103866001* from the TPSB subgroup; and *Thecc1EG029126t1*, *GSVIVT01016765001* and *GSVIVT01035467001* from TPSC subgroup) have been experimentally verified. In the TPSA subgroup, *GSVIVT01026868001* played an inhibitory role in flower development (Velasco et al., 2007), while both *Achn38246* and *Achn172901* acted as transcriptional activators involved in cold stress response (Park et al., 2010; Savage et al., 2013). Homologous genes in the TPSB subgroup, *Achn143561*, *Achn322351* and *GSVIVT0103866001*, performed a similar role in regulating plant protection against UV stress (Schenke et al., 2014). The paralogous genes of subgroup TPSC (*Thecc1EG029126t1*, *GSVIVT01016765001* and *GSVIVT01035467001*) mainly regulated plant epidermal cell fate (Cheng et al., 2014; Savage et al., 2013). Above all, we ventured that the function of the TPSA, TPSB and TPSC subgroups in *C. sinensis* might associate with responses to biotic and abiotic stress along with influencing certain developmental processes.

### Conserved motif analysis of *R2R3-MYB*s

The R2 and R3 MYB domains of the 118 *C. sinensis R2R3-MYB* TFs were analyzed (Fig. 3). The R2 and R3 domains contain a set of characteristic amino acids, which include the highly conserved and evenly distributed tryptophan residues (Trp, W) known to be critical for sequence-specific binding of DNA (Cao et al., 2013; Stracke et al., 2001), demonstrating that the R2 and R3 MYB repeats of the MYB DNA-binding domain are highly conserved in *C. sinensis*, consistent with previous findings of the counterpart genes in other plant lineages (Du et al., 2012; Li et al., 2016; Wilkins et al., 2009). Most of the conserved residues were situated between the second and third conserved W residues in each MYB repeat, elucidating that the first area of them was less conserved than the other two.

**Figure 3.**
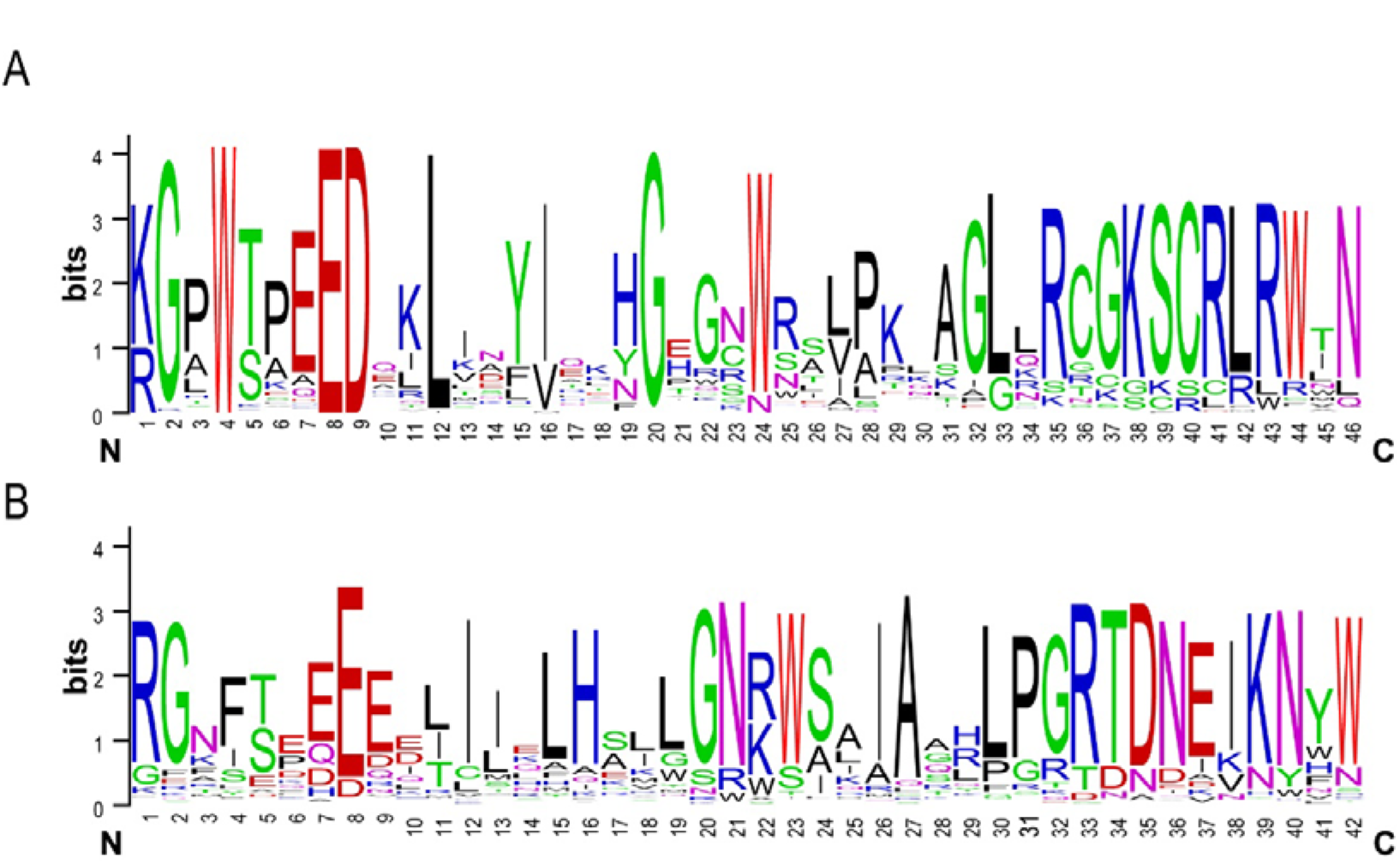
Analysis of R2 and R3 domains of *C. sinensis* R2R3-MYB TFs. The sequence logos of the R2 (A) and R3 (B) MYB repeats were determined via multiple sequence alignment of the R2R3-MYB proteins. The bit score indicates the information content for each position in the sequence. Highly conserved Trp residues critical for DNA binding in the MYB domain are highlighted in red.

Conserved amino acid motifs represent functional areas that are maintained during evolution. The conserved motifs within the 118 R2R3-MYB sequences were analyzed and aligned using MEME Suite (Bailey et al., 2009). A total of 20 conserved motifs were identified in the R2R3-MYB family (Fig. 4). Six of these, motifs 1 to 6, were present in all R2R3-MYB members except for CsMYB1a and thus were designated as “general motifs;” the rest of the motifs (motifs 7 to 20) were considered to be “specific motifs,” since they were present in only one or several R2R3-MYB members. For instance, motifs 16 and 9 were unique to CsMYB1a and CsMYB117a; meanwhile, motifs 15 and 19 were contained only in three genes. Overall, the members clustering to the same clade harbored similar motif patterns.

**Figure 4.**
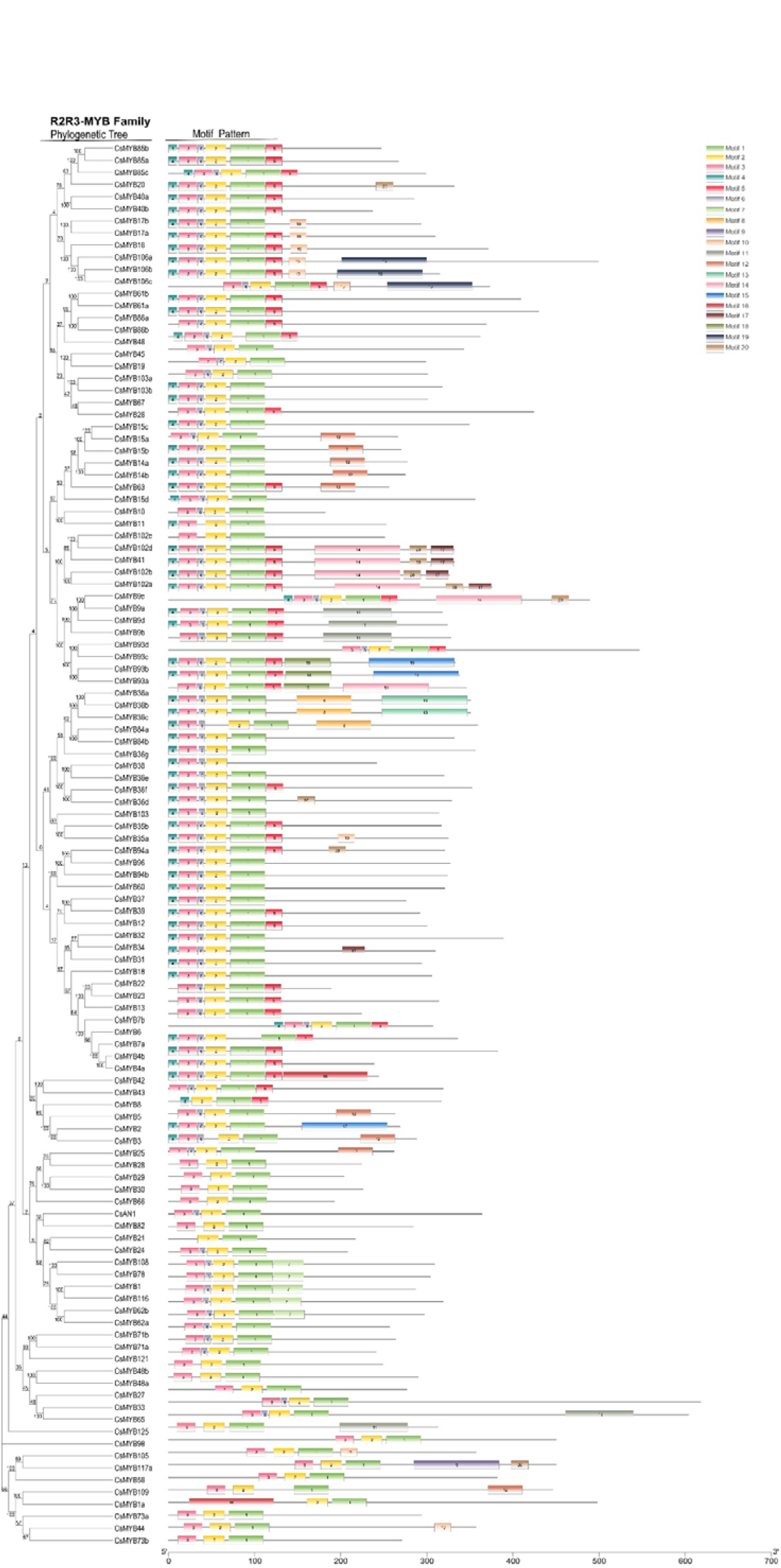
Phylogenetic relationships and conserved motifs of *C. sinensis* R2R3-MYB TFs. The neighbor-joining tree of 118 R2R3-MYB proteins is shown on the left, and the structures of 20 conserved motifs in R2R3-MYB TFs, predicted by MEME Suite, are shown on the right.

### Expression patterns of the *R2R3-MYB* family

The expression patterns of 118 genes encoding *R2R3-MYB* TFs were analyzed in different tissues. No transcripts were detected for *CsMYB101* (*TEA028392.1*) and *CsMYB117b* (*TEA002233.1*), suggesting they are pseudogenes in *C. sinensis*. The genes were classified into seven expression clusters, based on their distinct transcript patterns in various tissues and organs (Fig. 5). The 21 genes in RNA-seq-based cluster 1 were expressed mainly in flowers; genes in cluster 2 (14) were predominantly expressed in fruits; genes in cluster 3 (29) were mainly expressed in tender roots; genes in cluster 4 (8) were expressed at comparable levels in both apical buds and tender roots; cluster 5 genes (18) were mainly present in apical shoots and young leaves; while cluster 6 genes (21) were mainly expressed in stems and finally, cluster 7 genes (5) were equally expressed in mature leaves, old leaves and stems. Normally, genes within the same phylogenetic subgroup exhibit distinct transcript profiles (Dubos et al., 2010). Such was the case for subgroup S14: the members in this subgroup were detected in RNA-seq-based clusters 1, 3 and 6. In *A. thaliana*, members of S14 were generally related to axillary bud formation and cell differentiation. In some cases, however, genes belonging to the same subgroups might also have similar transcription profiles in the same tissue, organ or cell type. Such was the case with the TPSA subgroup; all were present in RNA-seq-based cluster 3. Likewise, all members of subgroup S5 gathered in cluster 5.

**Figure 5.**
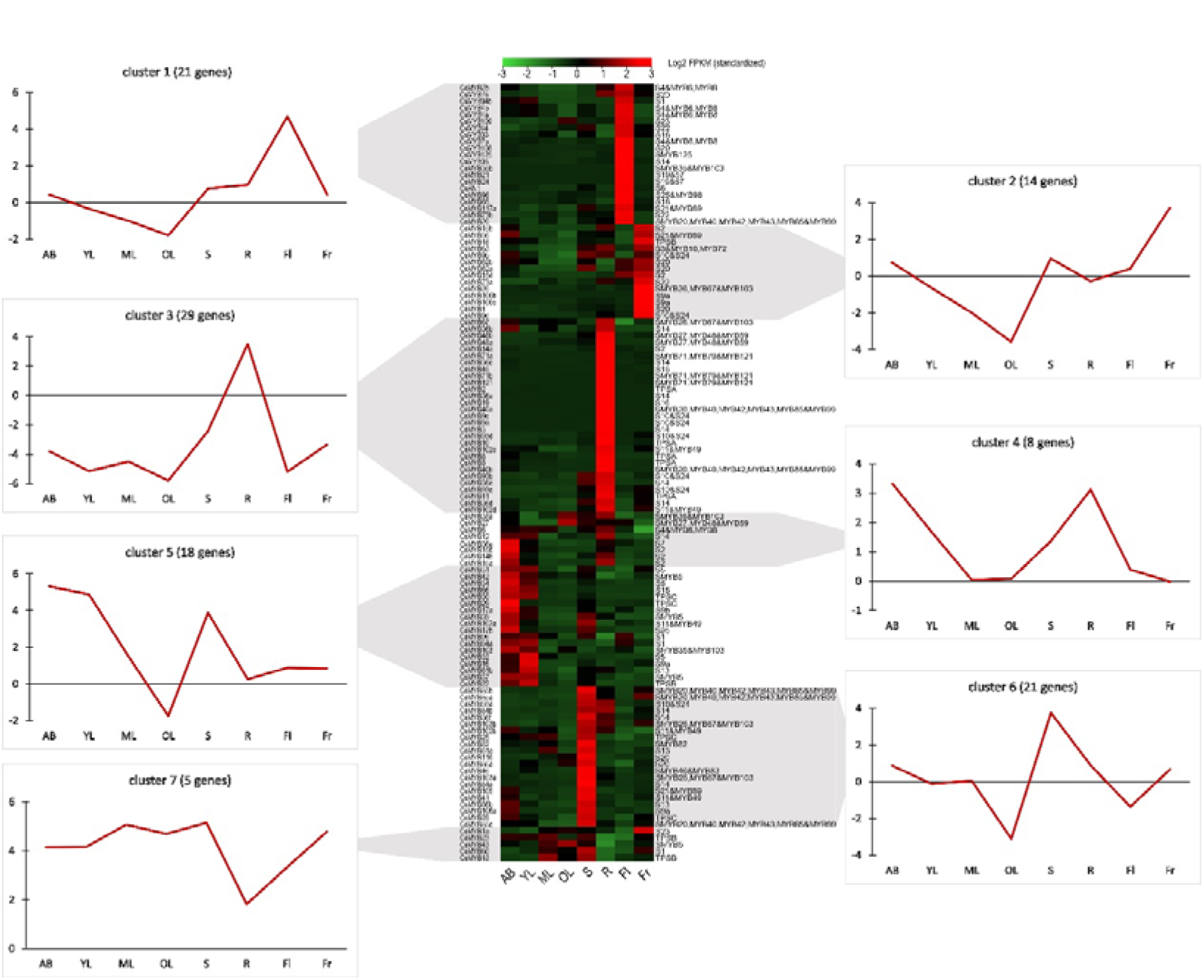
Heatmap of the 118 *CsR2R3-MYB* genes transcribed in the tissues of *C. sinensis*. The 118 *CsR2R3-MYB* genes clustered into seven expression groups, based on their tissue-specific expression. The gene name is indicated on the left of the heatmap, and the short name of the phylogenetic subgroup is on the right. Transcript abundance is expressed in standardized log2 fragments per kilobase of exon per million fragments mapped (FPKM) values. Next to each RNA-seq-based cluster, there is a graph with the mean transcript abundance for the entire cluster in each tissue. AB, apical bud; YL, young leaf; ML, mature leaf; OL, old leaf; S, stem; R, root; Fl, flower; Fr, fruit. Data were obtained from the Tea Plant Information Archive (http://tpia.teaplant.org/).

Previous reports have pointed out that most of the genes involved in flavonoid formation and catechin biosynthesis were preferentially expressed in apical buds and young leaves, where most galloylated catechins accumulate (Wei et al., 2018). Accordingly, *R2R3-MYB*s in RNA-seq-based cluster 5, whose expression level was highest in these two tender tissues, are the most likely candidates regulating catechin biosynthesis. As is shown in Fig. 5, almost half of the genes found in cluster 5 (9) belonged to the subgroups that specifically evolved in or expanded in *C. sinensis,* subgroups S5, SMYB5, TPSB and TPSC. It is worth noting that genes of subgroup S5 and SMYB5 were confirmed to be involved in flavonoid formation in *A. thaliana* (Nesi et al., 2001; Stracke et al., 2001), whereas subgroup TPSB was inferred to be relevant to UV protection and subgroup TPSC to control plant epidermal cell fate specification.

### *In silico* analysis of R2R3-MYB expression and catechin accumulation

Catechins are the major type of polyphenols, comprising up to 70% of the polyphenols in the tea plant. Fig 6B shows that catechin contents, especially the contents of the galloylated catechins EGCG and ECG, are significantly higher in apical buds and young leaves than in other tissues. Therefore, for understanding the possible correlation between the galloylated catechins and the *CsR2R3-MYB*s that specifically evolved or expanded in *C. sinensis*, we focused on the nine *R2R3-MYB*s grouped in cluster 5 (*CsMYB22*, *CsMYB29*, *CsMYB30*, *CsMYB31*, *CsMYB32*, *CsMYB34*, *CsMYB37*, *CsMYB39* and *CsMYB42*) because they were preferentially expressed in apical buds and young leaves.

**Figure 6.**
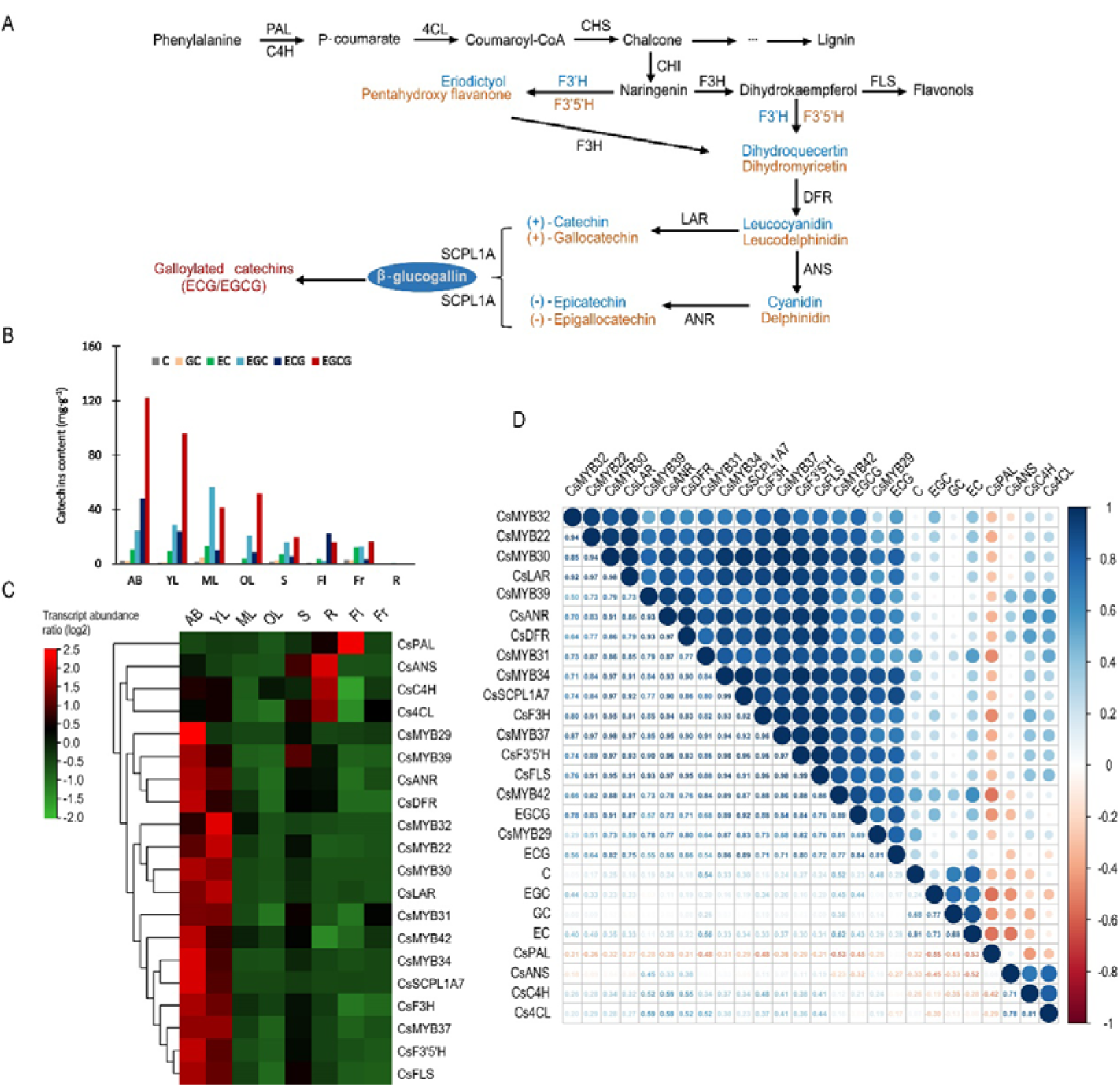
Expression pattern analysis and correlation analysis of tea-specific or tea-expanded *CsR2R3-MYB*s. (A) The biosynthetic pathway of catechins. *CHS*, *CHI*, *F3H*, *F3’H*, *F3’5’H*, *DFR*, *ANS*, *LAR*, *ANR* and *SCPL1A* represent genes encoding chalcone synthase, chalcone isomerase, flavanone 3-hydroxylase, flavonoid 3’-hydroxylase, flavonoid 3’,5’-hydroxylase, dihydroflavonol 4-reductase, anthocyanidin synthase, leucoanthocyanidin reductase, anthocyanidin reductase and type 1A serine carboxypeptidase-like acyltransferases, respectively. (B) The contents of six catechin monomers in eight tissues. (C) Heatmap of RNA-seq transcript abundance patterns of the 20 *CsR2R3-MYB* genes from the *C. sinensis* genome in eight different tissues. AB, apical bud; YL, young leaf; ML, mature leaf; OL, old leaf; S, stem; R, root; Fl, flower; Fr, fruit. (D) Correlative analysis of *CsR2R3-MYB* genes, structural genes and catechins accumulation patterns in eight representative tissues of *C. sinensis* plants. R>0.5 indicates a positive correlation; R<-0.5 indicates a negative correlation. Data were obtained from the Tea Plant Information Archive (http://tpia.teaplant.org/).

Further, the transcript abundance of the nine potential *R2R3-MYB* TFs and the catechin biosynthesis genes were investigated in different tissues. The results clearly showed that the genes in the catechin biosynthesis pathway display tissue-specific expression patterns. The genes downstream of catechin biosynthesis (*CsSCPL1A7*, *CsANR*, *CsLAR, CsDFR, CsFLS*, *CsF3H*, *CsF3’5’H)* were highly expressed in apical buds and young leaves, whereas the upstream genes (*CsPAL*, *Cs4CL, CsC4H*) were highly expressed in root and flower (Fig. 6C). Interestingly, the nine candidate *R2R3-MYB* TFs showed preferential expression in apical buds and young leaves, which was consistent with the expression pattern of the downstream catechin biosynthetic genes, indicating that those *R2R3-MYB*s have relevance to catechin biosynthesis.

To identify the relationship between transcript abundance and catechin contents, a comprehensive gene-to-metabolite correlation analysis was conducted. As shown in Fig. 6D, the three genes in the catechin biosynthesis pathway (*CsPAL*, *CsC4H* and *Cs4CL*) that were not tender parts-specific indeed showed a low correlation or negatively correlated to catechin content. Comparatively, the expression level of the nine candidate *R2R3-MYB* TFs was positively correlated to the transcript abundance patterns of the catechin biosynthesis pathway downstream genes (*CsSCPL1A7*, *CsANR*, *CsLAR, CsDFR, CsFLS*, *CsF3H*, *CsF3’5’H)* and correlated with the contents of EGCG and ECG, with inter-gene-to-metabolite Pearson’s correlation coefficients over 0.55. *CsMYB30*, *CsMYB34*, *CsMYB37* and *CsMYB42* exhibited good performance compared with the others, each having a correlation coefficient exceeding 0.75, indicating an extremely strong correlation. Notably, *CsMYB34* had the strongest correlation with the catechin biosynthesis pathway downstream genes (>0.9), especially with *CsSCPL1A7* (0.99). Besides that, the coefficients between *CsMYB34* and EGCG and ECG contents were 0.89 and 0.86, respectively.

### Validation of the correlation between *R2R3-MYB* TFs and catechins

To validate the relationship between *R2R3-MYB* TFs and catechins, HPLC and qRT-PCR assays were carried out. Different tissues of the tea plant were tested for the contents of different catechin monomers via HPLC (Fig. 7A). As shown in Fig. 7A, there was a high level of EGCG in all tested tissues compared with the other catechin monomers, reaching the highest level in apical buds (AB) and young leaves (FL and SL). Additionally, the expression patterns of the nine specially evolved and expanded candidate *CsR2R3-MYB*s and catechin biosynthesis genes in different tissues were verified by qRT-PCR. According to the relative expression patterns, there were four distinct clusters (cluster A-D) (Fig. 7B). The genes upstream of catechin biosynthesis (*CsPAL*, *CsC4H*, *Cs4CL*) grouped in cluster A and were highly expressed in roots, consistent with the absence of catechins in roots (Fig. 6C). *CsMYB22*, *CsMYB31* and *CsMYB42* (cluster B) had distinctly high expression in old leaves, and were not apical bud- or young leaf-specific. Remarkably, *CsMYB30*, *CsMYB32*, *CsMYB34* and *CsMYB37,* which clustered with critical genes downstream of catechins biosynthesis (cluster D) were highly expressed in apical buds and first leaves, where EGCG and ECG accumulated (Fig. 7A), preliminarily validating the intimate correlation between *CsMYB30*, *CsMYB32*, *CsMYB34*, *CsMYB37* and catechin biosynthesis.

**Figure 7.**
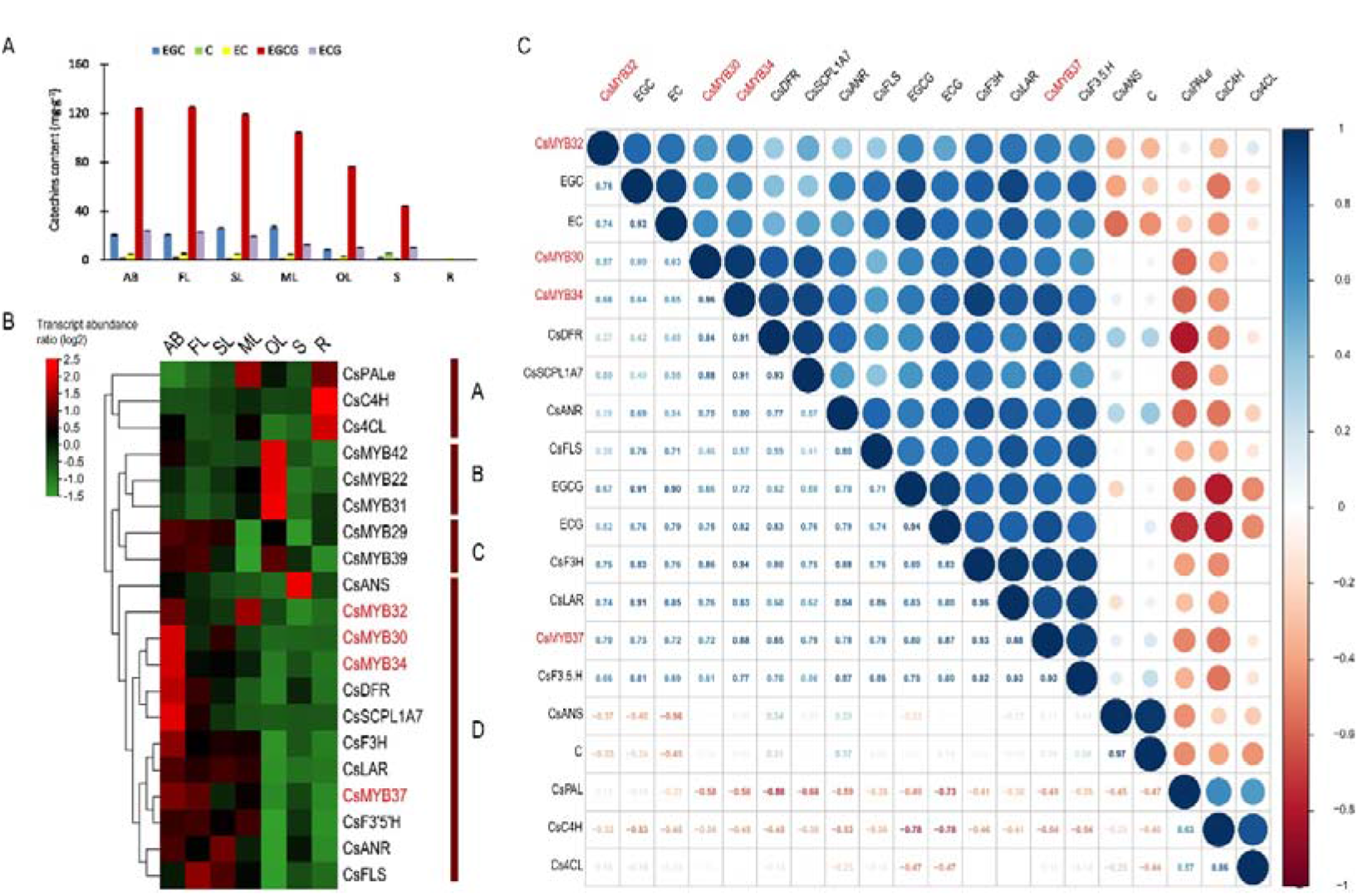
Expression profiling validation and correlation analysis. (A) The contents of six catechin monomers in seven tissues as measured with an HPLC system. (B) Heatmap of qRT-PCR transcript abundance patterns in seven different tissues. (C) Correlative analysis of four potential *CsR2R3-MYB* TFs, catechin biosynthesis genes and catechin accumulation patterns in different tissues of tea plants. R>0.5, positive correlation; R<-0.5, negative correlation. AB, apical bud; YL, young leaf; SL, second leaf; ML, mature leaf; OL, old leaf; S, stem; R, root.

For further confirmation, gene-to-metabolite correlation analysis of *CsMYB30*, *CsMYB32*, *CsMYB34*, *CsMYB37*, key genes of the catechin biosynthesis pathway, and the accumulation of catechins was conducted (Fig. 7C). The results confirmed the extremely low correlation between four of the candidate TFs and the genes upstream of catechin biosynthesis *CsPAL*, *CsC4H* and *Cs4CL*. In contrast, *CsMYB30*, *CsMYB34* and *CsMYB37* were strongly associated with most of the major genes downstream of the galloylated catechin biosynthesis pathway (*CsSCPL1A7*, *CsANR, CsLAR* and *CsDFR*) as well as ECG and EGCG contents (> 0.7). Particularly, *CsMYB37* showed the highest correlation level with the highest inter-correlation coefficients within the four major functional genes *CsSCPL1A7*, *CsANR*, *CsLAR, CsDFR* and the contents of EGCG and ECG reaching to 0.79, 0.78, 0.88, 0.85, 0.87 and 0.8, respectively. However, *CsMYB30*, *CsMYB34* and *CsMYB37* were less related to the other downstream catechin biosynthesis genes (*CsFLS*, *CsF3H*, *CsF3’5’H*) according to their lower Pearson’s correlation coefficients. Consistent with the *in silico* results in Fig. 6D, the correlation among *CsMYB32*, the galloylated catechin contents and the biosynthesis genes was weaker than the other three *R2R3-MYB*s. In more detail, it had a relatively low correlation coefficient with ECG (0.52) and EGCG (0.67), and a relatively low correlation coefficient with genes *CsSCPL1A, CsANR, CsLAR* and *CsDFR* (0.5, 0.36, 0.74 and 0.37, respectively).

The results provide convincing clues that *CsMYB30*, *CsMYB34*, *CsMYB37* might be the key transcription factors regulating galloylated catechins biosynthesis for the *Camellia*-specific specialized metabolites ECG and EGCG.

## Discussion

Plants are rich in metabolites that allow them to adapt to the environment and resist biotic and abiotic stress (Howe and Jander, 2008). These metabolites are widely used as natural products for treating human diseases and are valuable raw materials for modern industry (Guo et al., 2018; Fang et al., 2019; Howe et al., 2008; Plomion et al., 2001). *C. sinensis* is an advantageous model system to dig into plant-specific metabolite biosynthesis and genetic regulation. Its metabolites, such as flavonoids, caffeine, and volatile terpenes, accumulate in abundance and share characteristics with the same metabolites in other plant lineages (Yu et al., 2020).

Galloylated catechins are secondary metabolites only found in *Vitis vinifera* and *C. sinensis* (Wei et al., 2018). Contrary to the small amount of galloylated catechins present in the form of ECG in *Vitis vinifera* (Bontpart et al., 2018), they are abundant in *C. sinensis*, and EGCG, only existing in tea plants, is the predominant form (Kim et al., 2004; Steinmann et al., 2013). It is the most bioactive component among the catechin enantiomers, and is derived from the flavonoid branch of the phenylpropanoid metabolite pathway. The acyltransferase family, belonging to subclade 1A of serine carboxypeptidase-like (SCPL) acyltransferases, acts as the most critical downstream gene family involved in the production of EGCG and ECG (Wei et al., 2018). This family extensively expanded to 22 members in the *C. sinensis* genome, while the *Vitis vinifera* genome contains half that number (11) (Wei et al., 2018). Two key enzymes Epicatechin:1-O-galloyl-b-D-glucose O-galloyltransferase (ECGT) and UDP-glucose: galloyl-1-O-b-D-glucosyltransferase (UGGT) are recruited to catalyze the last two reactions in this bioprocess (Liu et al., 2012). The biosynthesis of galloylated catechins in *C. sinensis* has been comprehensively investigated with regard to the biosynthesis genes of the pathway. However, the transcriptional regulation of these pathways remains to be illuminated.

Genes responsible for plant secondary metabolite biosynthesis are coordinately regulated by TFs, a regulatory superfamily that dynamically drives the evolution of plant metabolic pathways for special compounds (Shoji et al., 2021). The regulatory network of this gene superfamily is highly conserved both in angiosperms and gymnosperms (Zhang et al., 2014). The *R2R3-MYB* TFs confer tissue-specific or development stage-specific patterns for metabolites in the same biosynthesis pathway; often, multiple paralogues coexist in one species (Zhang et al., 2014). Lignin, flavonoids, anthocyanins and capsaicinoids are four different types of secondary metabolites synthesized from the phenylpropanoid pathway that are regulated by *R2R3-MYB* TFs (Sun et al., 2016; Zhu et al., 2019; Soler et al., 2015). Remarkably, the great expansion of this transcription-regulatory superfamily in plant lineages appears to account for the diversity of regulatory functions that the *R2R3-MYB* TFs undertake in plant-specific metabolic bioprocesses (Millard et al., 2019). As demonstrated in detail by the analysis of Soler et.al, the R2R3-MYB subgroups in *E. grandis*, *V. vinifera* and *P. trichocarpa*, which were equipped with expanded members, greatly determined the diversification of specific functions in lignin biosynthesis (Soler et al., 2015).

Based on the consideration of the unique and abundant accumulation of the specific galloylated catechins (ECG and EGCG) in tea plant and tea-specific *R2R3-MYB* TFs identified in this work, we hypothesize that the biosynthesis of the characteristic galloylated catechins is absolutely influenced at the transcription regulation level. Different from the result that concentrates on the involvement of some *CsR2R3-MYB* genes in response to drought, cold, gibberellic acid (GA), and abscisic acid (ABA) treatments, which are revealed in the recent genome-wide report of this family(Chen et al., 2021), we firstly emphasize on confirming the putative *R2R3-MYB* candidates that directly function in the production of *Camellia*-specialized compounds (EGCG) among the tea-specific *R2R3-MYB* transcription factors.

The comprehensive and comparative phylogenetic analysis of *CsR2R3-MYB* TFs, backed by multiple sequence alignment among *C. sinensis* and *A. thaliana,* suggests that most of the members in this family are conserved. Most *R2R3-MYB*s share similar functions to the homologous counterparts studied in *A. thaliana*. Some of the *R2R3-MYB* TFs that clustered in TPSA, TPSB and TPSC subgroups evolved exclusively in *C. sinensis*, but have isogenous genes in *Actinidia chinensis*, *Vitis vinifera*, *Theobroma cacao* and *Coffea canephora* genomes. Thus, we speculated that TPSA, TPSB and TPSC are either obtained in *C. sinensis* or lost in *A. thaliana* lineages after divergence from their most recent common ancestor during two whole-genome duplication (WGD) events. In addition, members of SMYB5 and S5 subgroups, regulating flavonoids biosynthesis in *A. thaliana*, are greatly expanded in *C. sinensis*, which suggests that they might be either functionally redundant genes or genes that undertake some novel functions in the tea plant.

Considering that characteristic catechins highly accumulate in apical buds and young leaves, we speculated that the *CsR2R3-MYB* TFs that are preferentially expressed in these tender tissues along with the major catechin-biosynthesis genes are the most promising candidates putatively regulating the biosynthesis of tea-specific catechins. Consistent with previous results (Wei et al., 2018), our study observed high expression levels of key galloylated catechin biosynthesis genes *SCPL1A, ANR, LAR* and *DFR* in tender tissues, while the expression of upstream genes (*PAL*, *C4H*, *4CL*) in the phenylpropanoid pathway that are mainly relevant to the generation of condensed polymer proanthocyanidins (PAs), was in fruits, flowers and roots. However, *CsMYB42* was preferentially expressed in tender tissues and had a strong correlation with catechin biosynthesis genes and the contents of ECG and EGCG in the ‘Shuchazao’ variety (Fig. 6D), while it was preferentially expressed in old leaves in the ‘Lingtoudancong’ variety (Fig. 7C). Thus, differences can be observed in different *C. sinensis* varieties. Eventually, through systematic analyses, *CsMYB30* (TPSC subgroup), *CsMYB34* (S5 subgroup) and *CsMYB37* (SMYB5 subgroup) were confirmed as the potential *R2R3-MYB* TFs relevant to the internal accumulation of characteristic catechins (ECG and EGCG) in *C. sinensis*, however, further investigation is needed. This study laid a theoretical framework and valuable foundation for the needed future work, as we have provided a considerable amount of preliminarily evidence through systematic bioinformatics analysis to gain a deeper perception of the functional roles of the R2R3-MYB superfamily in *C. sinensis*. Nevertheless, it is still necessary to further exploration and validate these results.

## Conclusions

A total of 118 *R2R3-MYB* gene members, classified into 38 subgroups, were identified in the *C. sinensis* genome. Notably, five subgroups (TPSA, TPSB, TPSC, S5 and SMYB5) containing 21 *R2R3-MYB* TFs were identified to be remarkably expanded in or completely unique to *C. sinensis*. Furthermore, gene structure predictions, expression profile validation and correlation analyses were subsequently conducted to screen out the most promising candidate *R2R3-MYB* TFs (*CsMYB30*, *CsMYB34* and *CsMYB37*) that positively function in galloylated catechin biosynthesis in tea plants. The present findings underpin a basic understanding of species-specific regulatory mechanisms that *C. sinensis* employs to biosynthesize specialized metabolites and will be beneficial for selecting favorable *C. sinensis* germplasms.

## Abbreviation

C: catechin
EC: epicatechin
GC: gallatechin
EGC: epigallocatechin
CG: catechin gallate
ECG: epigallocatechin gallate
GCG: gallatechin gallate
EGCG: epigallocatechin gallate
qRT-PCR: Quantitative reverse transcription polymerase chain reaction
PAL: Phenylalanine ammonia lyase
C4H: Cinnamate 4-hydroxylase
4CL: 4-Coumarate: coenzyme A ligase
CHS: Chalcone synthase
CHI: Chalcone isomerase
F3H: Flavanone -3-hydroxylase
F3’H: Flavonoid 3’-hydroxylase
F3’5’H: Flavonoid 3’5’-hydroxylase
FLS: Flavonol synthase
DFR: Dihydroflavonol 4-reductase
ANS: Anthocyanidin synthase
ANR: Anthocyanidin reductase
LAR: Leucoanthcyanidin 4-reductase
SCPL1A: Subclade 1A of serine carboxypeptidase-like acyltransferases
TFs: transcription factors
HPLC: high-performance liquid chromatography
Trp/W: tryptophan

## Data availablity statement

The original contributions presented in the study are included in the article/Supplementary Material, further inquiries can be directed to the corresponding author/s.

## Authors’ contributions

J.L. performed the qRT-PCR test, analyzed the data, made the data charts and wrote the manuscript. S.L. conceived the project and supervised the researches. P.C. carried out the HPLC test. J.C., S.T. and W.Y collected the materials for the experiments and provided useful suggestions. F.C. reviewed and edited the manuscript. P.Z funded the researches and reviewed the manuscript. B.S. designed the project, supervised the researches, interpreted data and edited the manuscript.

## Appendix A. Supplementary data

## References

1. Abbas, M., Saeed, F., Anjum, F.M., Afzaal, M., Tufail, T., Bashir, M.S., Ishtiaq, A., Hussain, S., Suleria, H.A.R., 2017. Natural polyphenols: An overview. Int. J. Food Prop. 20, 1689–1699. https://doi.org/10.1080/10942912.2016.1220393

2. Ahmad, N., Feyes, D.K., Agarwal, R., Mukhtar, H., Nieminen, A.-L., 1997. Green tea constituent epigallocatechin-3-gallate and induction of apoptosis and cell cycle arrest in human carcinoma cells. JNCI J. Natl. Cancer Inst. 89, 1881–1886. https://doi.org/10.1093/jnci/89.24.1881

3. Ahmad, N., Gupta, S., Mukhtar, H., 2000. Green tea polyphenol epigallocatechin-3-gallate differentially modulates nuclear factor κB in cancer cells versus normal cells. Arch. Biochem. Biophys. 376, 338–346. https://doi.org/10.1006/abbi.2000.1742

4. Ambawat, S., Sharma, P., Yadav, N.R., Yadav, R.C., 2013. MYB transcription factor genes as regulators for plant responses: an overview. Physiol. Mol. Biol. Plants 19, 307–321. https://doi.org/10.1007/s12298-013-0179-1

5. Asakawa, T., Hamashima, Y., Kan, T., 2013. Chemical synthesis of tea polyphenols and related compounds. Curr. Pharm. Des. 19, 6207–6217. https://doi.org/10.2174/1381612811319340012

6. Ashihara, H., Deng, W.-W., Mullen, W., Crozier, A., 2010. Distribution and biosynthesis of flavan-3-ols in *Camellia sinensis* seedlings and expression of genes encoding biosynthetic enzymes. Phytochemistry 71, 559–566. https://doi.org/https://doi.org/10.1016/j.phytochem.2010.01.010

7. Bailey, T.L., Boden, M., Buske, F.A., Frith, M., Grant, C.E., Clementi, L., Ren, J., Li, W.W., Noble, W.S., 2009. MEME Suite: Tools for motif discovery and searching. Nucleic Acids Res. 37, 202–208. https://doi.org/10.1093/nar/gkp335

8. Bedon, F., Grima-Pettenati, J., Mackay, J., 2007. Conifer R2R3-MYB transcription factors: sequence analyses and gene expression in wood-forming tissues of white spruce (Picea glauca). BMC Plant Biol. 7, 17. https://doi.org/10.1186/1471-2229-7-17

9. Bontpart, T., Ferrero, M., Khater, F., Marlin, T., Vialet, S., Vallverdù-Queralt, A., Pinasseau, L., Ageorges, A., Cheynier, V., Terrier, N., 2018. Focus on putative serine carboxypeptidase-like acyltransferases in grapevine. Plant Physiol. Biochem. 130, 356–366. https://doi.org/10.1016/j.plaphy.2018.07.023

10. Cao, Z.-H., Zhang, S.-Z., Wang, R.-K., Zhang, R.-F., Hao, Y.-J., 2013. Genome wide analysis of the apple MYB transcription factor family allows the identification of *MdoMYB121* gene confering abiotic stress tolerance in plants. PLoS One 8, e69955. https://doi.org/10.1371/journal.pone.0069955

11. Chen, C., Chen, H., Zhang, Y., Thomas, H.R., Frank, M.H., He, Y., Xia, R., 2020. TBtools: An integrative toolkit developed for interactive analyses of big biological data. Mol. Plant 13, 1194–1202. https://doi.org/10.1016/j.molp.2020.06.009

12. Chen, X., Wang, P., Gu, M., Lin, X., Hou, B., Zheng, Y., Sun, Y., Jin, S., Ye, N., 2021. R2R3-MYB transcription factor family in tea plant (*Camellia sinensis*): Genome-wide characterization, phylogeny, chromosome location, structure and expression patterns. Genomics 113, 1565–1578. https://doi.org/10.1016/j.ygeno.2021.03.033

13. Chen, Y., Yu, M., Xu, J., Chen, X., Shi, J., 2009. Differentiation of eight tea (*Camellia sinensis*) cultivars in China by elemental fingerprint of their leaves. J. Sci. Food Agric. 89, 2350–2355. https://doi.org/10.1002/jsfa.3716

14. Cheng, Y., Zhu, W., Chen, Y., Ito, S., Asami, T., Wang, X., 2014. Brassinosteroids control root epidermal cell fate via direct regulation of a MYB-bHLH-WD40 complex by GSK3-like kinases. Elife 3. https://doi.org/10.7554/eLife.02525

15. Drew, B., 2019. The growth of tea. Nature 566, S2–S4. https://doi.org/10.1038/d41586-019-00395-4

16. Du, H., Yang, S.-S., Liang, Z., Feng, B.-R., Liu, L., Huang, Y.-B., Tang, Y.-X., 2012. Genome-wide analysis of the MYB transcription factor superfamily in soybean. BMC Plant Biol. 12, 106. https://doi.org/10.1186/1471-2229-12-106

17. Dubos, C., Stracke, R., Grotewold, E., Weisshaar, B., Martin, C., Lepiniec, L., 2010. MYB transcription factors in *Arabidopsis*. Trends Plant Sci. 15, 573–581. https://doi.org/10.1016/j.tplants.2010.06.005

18. Eungwanichayapant, P.D., Popluechai, S., 2009. Accumulation of catechins in tea in relation to accumulation of mRNA from genes involved in catechin biosynthesis. Plant Physiol. Biochem. 47, 94–97. https://doi.org/https://doi.org/10.1016/j.plaphy.2008.11.002

19. Fang, C., Fernie, A.R., Luo, J., 2019. Exploring the diversity of plant metabolism. Trends Plant Sci. 24, 83–98. https://doi.org/https://doi.org/10.1016/j.tplants.2018.09.006

20. Fukai, K., Ishigami, T., Hara, Y., 1991. Antibacterial activity of tea polyphenols against phytopathogenic bacteria. Agric. Biol. Chem. 55, 1895–1897. https://doi.org/10.1080/00021369.1991.10870886

21. Gigolashvili, T., Yatusevich, R., Berger, B., Müller, C., Flügge, U.-I., 2007. The R2R3-MYB transcription factor *HAG1/MYB28* is a regulator of methionine-derived glucosinolate biosynthesis in *Arabidopsis thaliana*. Plant J. 51, 247–261. https://doi.org/10.1111/j.1365-313X.2007.03133.x

22. Goicoechea, M., Lacombe, E., Legay, S., Mihaljevic, S., Rech, P., Jauneau, A., Lapierre, C., Pollet, B., Verhaegen, D., Chaubet-Gigot, N., Grima-Pettenati, J., 2005. *EgMYB2*, a new transcriptional activator from Eucalyptus xylem, regulates secondary cell wall formation and lignin biosynthesis. Plant J. 43, 553–567. https://doi.org/https://doi.org/10.1111/j.1365-313X.2005.02480.x

23. Gonzalez, A., Mendenhall, J., Huo, Y., Lloyd, A., 2009. *TTG1* complex MYBs, *MYB5* and *TT2*, control outer seed coat differentiation. Dev. Biol. 325, 412–421. https://doi.org/https://doi.org/10.1016/j.ydbio.2008.10.005

24. Guo, L., Winzer, T., Yang, X., Li, Y., Ning, Z., He, Z., Teodor, R., Lu, Y., Bowser, T.A., Graham, I.A., Ye, K., 2018. The opium poppy genome and morphinan production. Science 362, 343–347. https://doi.org/10.1126/science.aat4096

25. He, X., Zhao, X., Gao, L., Shi, X., Dai, X., Liu, Y., Xia, T., Wang, Y., 2018. Isolation and characterization of key genes that promote flavonoid accumulation in purple-leaf tea (C*amellia sinensis* L.). Sci. Rep. 8, 1–13. https://doi.org/10.1038/s41598-017-18133-z

26. Heim, K.E., Tagliaferro, A.R., Bobilya, D.J., 2002. Flavonoid antioxidants: chemistry, metabolism and structure-activity relationships. J. Nutr. Biochem. 13, 572–584. https://doi.org/10.1016/s0955-2863(02)00208-5

27. Hichri, I., Barrieu, F., Bogs, J., Kappel, C., Delrot, S., Lauvergeat, V., 2011. Recent advances in the transcriptional regulation of the flavonoid biosynthetic pathway. J. Exp. Bot. 62, 2465–2483. https://doi.org/10.1093/jxb/erq442

28. Howe, G.A., Jander, G., 2008. Plant immunity to insect herbivores. Annu. Rev. Plant Biol. 59, 41–66. https://doi.org/10.1146/annurev.arplant.59.032607.092825

29. Jiang, C., Gu, X., Peterson, T., 2004. Identification of conserved gene structures and carboxy-terminal motifs in the Myb gene family of *Arabidopsis* and *Oryza sativa* L. ssp. *indica*. Genome Biol. 5, R46. https://doi.org/10.1186/gb-2004-5-7-r46

30. Jin, H., Martin, C., 1999. Multifunctionality and diversity within the plant MYB-gene family. Plant Mol. Biol. 41, 577–585. https://doi.org/10.1023/a:1006319732410

31. Jiu, S., Guan, L., Leng, X., Zhang, K., Haider, M.S., Yu, X., Zhu, X., Zheng, T., Ge, M., Wang, C., Jia, H., Shangguan, L., Zhang, C., Tang, X., Abdullah, M., Javed, H.U., Han, J., Dong, Z., Fang, J., 2021. The role of *VvMYBA2r* and *VvMYBA2w* alleles of the *MYBA2* locus in the regulation of anthocyanin biosynthesis for molecular breeding of grape (*Vitis* spp.) skin coloration. Plant Biotechnol. J. 19, 1216–1239. https://doi.org/10.1111/pbi.13543

32. Kim, S.Y., Ahn, B.H., Min, K.J., Lee, Y.H., Joe, E.H., Min, D.S., 2004. Phospholipase D isozymes mediate epigallocatechin gallate-induced cyclooxygenase-2 expression in astrocyte cells. J. Biol. Chem. 279, 38125. https://doi.org/10.1074/jbc.M402085200

33. Kranz, H.D., Denekamp, M., Greco, R., Jin, H., Leyva, A., Meissner, R.C., Petroni, K., Urzainqui, A., Bevan, M., Martin, C., Smeekens, S., Tonelli, C., Paz-Ares, J., Weisshaar, B., 1998. Towards functional characterisation of the members of the *R2R3-MYB* gene family from *Arabidopsis thaliana*. Plant J. 16, 263–276. https://doi.org/10.1046/j.1365-313x.1998.00278.x

34. Kumar, S., Stecher, G., Tamura, K., 2016. MEGA7: Molecular evolutionary genetics analysis version 7.0 for bigger datasets. Mol. Biol. Evol. 33, 1870–1874. https://doi.org/10.1093/molbev/msw054

35. Li, M., Li, Y., Guo, L., Gong, N., Pang, Y., Jiang, W., Liu, Y., Jiang, X., Zhao, L., Wang, Y., Xie, D.-Y., Gao, L., Xia, T., 2017. Functional characterization of tea (*Camellia sinensis*) *MYB4a* transcription factor using an integrative approach. Front. Plant Sci. 8, 943. https://doi.org/10.3389/fpls.2017.00943

36. Li, S., Sun, L., Fan, X., Zhang, Y., Jiang, J., Liu, C., 2020. Functional analysis of *Vitis davidii* R2R3-MYB transcription factor *VdMYB14* in the regulation of flavonoid biosynthesis. J. Fruit Sci. 37, 783–792. https://doi.org/10.13925/j.cnki.gsxb.20190577

37. Li, W., Ding, Z., Ruan, M., Yu, X., Peng, M., Liu, Y., 2017. Kiwifruit R2R3-MYB transcription factors and contribution of the novel *AcMYB75* to red kiwifruit anthocyanin biosynthesis. Sci. Rep. 7, 16861. https://doi.org/10.1038/s41598-017-16905-1

38. Li, X., Xue, C., Li, J., Qiao, X., Li, L., Yu, L., Huang, Y., Wu, J., 2016. Genome-wide identification, evolution and functional divergence of MYB transcription factors in Chinese White Pear (*Pyrus bretschneideri*). Plant Cell Physiol. 57, 824–847. https://doi.org/10.1093/pcp/pcw029

39. Lipsick, J.S., 1996. One billion years of Myb. Oncogene 13, 223–235.

40. Liu, Y., Gao, L., Liu, L., Yang, Q., Lu, Z., Nie, Z., Wang, Y., Xia, T., 2012. Purification and characterization of a novel galloyltransferase involved in catechin galloylation in the tea plant (*Camellia sinensis*)*. J. Biol. Chem. 287, 44406–44417. https://doi.org/https://doi.org/10.1074/jbc.M112.403071

41. Livak, K.J., Schmittgen, T.D., 2001. Analysis of relative gene expression data using real-time quantitative PCR and the 2(-Delta Delta C(T)) Method. Methods 25, 402–408. https://doi.org/10.1006/meth.2001.1262

42. Martin, C., Paz-Ares, J., 1997. MYB transcription factors in plants. Trends Genet. 13, 67–73. https://doi.org/https://doi.org/10.1016/S0168-9525(96)10049-4

43. Matus, J.T., Aquea, F., Arce-Johnson, P., 2008. Analysis of the grape MYB R2R3 subfamily reveals expanded wine quality-related clades and conserved gene structure organization across *Vitis* and *Arabidopsis* genomes. BMC Plant Biol. 8, 83. https://doi.org/10.1186/1471-2229-8-83

44. Mehrtens, F., Kranz, H., Bednarek, P., Weisshaar, B., 2005. The *Arabidopsis* transcription factor MYB12 is a flavonol-specific regulator of phenylpropanoid biosynthesis. Plant Physiol. 138, 1083–1096. https://doi.org/10.1104/pp.104.058032

45. Millard, P.S., Kragelund, B.B., Burow, M., 2019. R2R3 MYB transcription factors – functions outside the DNA-binding domain. Trends Plant Sci. 24, 934–946. https://doi.org/https://doi.org/10.1016/j.tplants.2019.07.003

46. Mondal, T.K., Bhattacharya, A., Laxmikumaran, M., Ahuja, P.S., 2004. Recent advances of tea (*Camellia sinensis*) biotechnology. Plant Cell. Tissue Organ Cult. 76, 195–254. https://doi.org/10.1023/B:TICU.0000009254.87882.71

47. Nesi, N., Jond, C., Debeaujon, I., Caboche, M., Lepiniec, L., 2001. The *Arabidopsis TT2* gene encodes an R2R3 MYB domain protein that acts as a key determinant for proanthocyanidin accumulation in developing seed. Plant Cell 13, 2099–2114. https://doi.org/10.1105/tpc.010098

48. Nikoo, M., Regenstein, J.M., Ahmadi Gavlighi, H., 2018. Antioxidant and antimicrobial activities of (-)-epigallocatechin-3-gallate (EGCG) and its potential to preserve the quality and safety of foods. Compr. Rev. Food Sci. Food Saf. 17, 732–753. https://doi.org/10.1111/1541-4337.12346

49. Ogata, K., Morikawa, S., Nakamura, H., Hojo, H., Yoshimura, S., Zhang, R., Aimoto, S., Ametani, Y., Hirata, Z., Sarai, A., Ishii, S., Nishimura, Y., 1995. Comparison of the free and DNA-complexed forms of the DMA-binding domain from c-Myb. Nat. Struct. Biol. 2, 309–320. https://doi.org/10.1038/nsb0495-309

50. Park, M.-R., Yun, K.-Y., Mohanty, B., Herath, V., Xu, F., Wijaya, E., Bajic, V.B., Yun, S.-J., De Los Reyes, B.G., 2010. Supra-optimal expression of the cold-regulated OsMyb4 transcription factor in transgenic rice changes the complexity of transcriptional network with major effects on stress tolerance and panicle development. Plant. Cell Environ. 33, 2209–2230. https://doi.org/10.1111/j.1365-3040.2010.02221.x

51. Patzlaff, A., McInnis, S., Courtenay, A., Surman, C., Newman, L.J., Smith, C., Bevan, M.W., Mansfield, S., Whetten, R.W., Sederoff, R.R., Campbell, M.M., 2003. Characterisation of a pine MYB that regulates lignification. Plant J. 36, 743–754. https://doi.org/https://doi.org/10.1046/j.1365-313X.2003.01916.x

52. Plomion, C., Leprovost, G., Stokes, A., 2001. Wood formation in trees. Plant Physiol. 127, 1513–1523. https://doi.org/10.1104/pp.010816

53. Punyasiri, P.A.N., Abeysinghe, I.S.B., Kumar, V., Treutter, D., Duy, D., Gosch, C., Martens, S., Forkmann, G., Fischer, T.C., 2004. Flavonoid biosynthesis in the tea plant *Camellia sinensis*: properties of enzymes of the prominent epicatechin and catechin pathways. Arch. Biochem. Biophys. 431, 22–30. https://doi.org/https://doi.org/10.1016/j.abb.2004.08.003

54. Sakanaka, S., Juneja, L.R., Taniguchi, M., 2000. Antimicrobial effects of green tea polyphenols on thermophilic spore-forming bacteria. J. Biosci. Bioeng. 90, 81–85. https://doi.org/10.1016/s1389-1723(00)80038-9

55. Savage, N., Yang, T.J.W., Chen, C.Y., Lin, K.-L., Monk, N.A.M., Schmidt, W., 2013. Positional signaling and expression of *ENHANCER OF TRY AND CPC1* are tuned to increase root hair density in response to phosphate deficiency in *Arabidopsis thaliana*. PLoS One 8, e75452. https://doi.org/10.1371/journal.pone.0075452

56. Schenke, D., Cai, D., Scheel, D., 2014. Suppression of UV-B stress responses by flg22 is regulated at the chromatin level via histone modification. Plant. Cell Environ. 37, 1716–1721. https://doi.org/10.1111/pce.12283

57. Shoji, T., Umemoto, N., Saito, K., 2021. Genetic divergence in transcriptional regulators of defense metabolism: insight into plant domestication and improvement. Plant Mol. Biol. https://doi.org/10.1007/s11103-021-01159-3

58. Singh, K., Kumar, S., Rani, A., Gulati, A., Ahuja, P.S., 2009a. Phenylalanine ammonia-lyase (PAL) and cinnamate 4-hydroxylase (C4H) and catechins (flavan-3-ols) accumulation in tea. Funct. Integr. Genomics 9, 125–134. https://doi.org/10.1007/s10142-008-0092-9

59. Singh, K., Rani, A., Kumar, S., Sood, P., Mahajan, M., Yadav, S.K., Singh, B., Ahuja, P.S., 2008. An early gene of the flavonoid pathway, flavanone 3-hydroxylase, exhibits a positive relationship with the concentration of catechins in tea (*Camellia sinensis*). Tree Physiol. 28, 1349–1356. https://doi.org/10.1093/treephys/28.9.1349

60. Singh, K., Rani, A., Paul, A., Dutt, S., Joshi, R., Gulati, A., Ahuja, P.S., Kumar, S., 2009b. Differential display mediated cloning of anthocyanidin reductase gene from tea (*Camellia sinensis*) and its relationship with the concentration of epicatechins. Tree Physiol. 29, 837–846. https://doi.org/10.1093/treephys/tpp022

61. Soler, M., Camargo, E.L.O., Carocha, V., Cassan-Wang, H., San Clemente, H., Savelli, B., Hefer, C.A., Paiva, J.A.P., Myburg, A.A., Grima-Pettenati, J., 2015. The eucalyptus grandis R2R3-MYB transcription factor family: Evidence for woody growth-related evolution and function. New Phytol. 206, 1364–1377. https://doi.org/10.1111/nph.13039

62. Steinmann, J., Buer, J., Pietschmann, T., Steinmann, E., 2013. Anti-infective properties of epigallocatechin-3-gallate (EGCG), a component of green tea. Br. J. Pharmacol. 168, 1059–1073. https://doi.org/10.1111/bph.12009

63. Stracke, R., Werber, M., Weisshaar, B., 2001. The R2R3-MYB gene family in *Arabidopsis thaliana*. Curr. Opin. Plant Biol. 4, 447–456. https://doi.org/10.1016/s1369-5266(00)00199-0

64. Sun, B., Zhu, Z., Cao, P., Chen, H., Chen, C., Zhou, X., Mao, Y., Lei, J., Jiang, Y., Meng, W., Wang, Y., Liu, S., 2016. Purple foliage coloration in tea (*Camellia sinensis* L.) arises from activation of the R2R3-MYB transcription factor CsAN1. Sci. Rep. 6. https://doi.org/10.1038/srep32534

65. Surh, Y.-J., 2003. Cancer chemoprevention with dietary phytochemicals. Nat. Rev. Cancer 3, 768–780. https://doi.org/10.1038/nrc1189

66. Taguri, T., Tanaka, T., Kouno, I., 2004. Antimicrobial activity of 10 different plant polyphenols against bacteria causing food-borne disease. Biol. Pharm. Bull. 27, 1965–1969. https://doi.org/10.1248/bpb.27.1965

67. Teng, S., Keurentjes, J., Bentsink, L., Koornneef, M., Smeekens, S., 2005. Sucrose-specific induction of anthocyanin biosynthesis in *Arabidopsis* requires the *MYB75/PAP1* gene. Plant Physiol. 139, 1840–1852. https://doi.org/10.1104/pp.105.066688

68. Velasco, R., Zharkikh, A., Troggio, M., Cartwright, D.A., Cestaro, A., Pruss, D., Pindo, M., FitzGerald, L.M., Vezzulli, S., Reid, J., Malacarne, G., Iliev, D., Coppola, G., Wardell, B., Micheletti, D., Macalma, T., Facci, M., Mitchell, J.T., Perazzolli, M., Eldredge, G., Gatto, P., Oyzerski, R., Moretto, M., Gutin, N., Stefanini, M., Chen, Y., Segala, C., Davenport, C., Dematté, L., Mraz, A., Battilana, J., Stormo, K., Costa, F., Tao, Q., Si-Ammour, A., Harkins, T., Lackey, A., Perbost, C., Taillon, B., Stella, A., Solovyev, V., Fawcett, J.A., Sterck, L., Vandepoele, K., Grando, S.M., Toppo, S., Moser, C., Lanchbury, J., Bogden, R., Skolnick, M., Sgaremella, V., Bhatnagar, S.K., Fontana, P., Gutin, A., Van de Peer, Y., Salamini, F., Viola, R., 2007. A high quality draft consensus sequence of the genome of a heterozygous grapevine variety. PLoS One 2. https://doi.org/10.1371/journal.pone.0001326

69. Wei, C., Yang, H., Wang, S., Zhao, J., Liu, C., Gao, L., Xia, E., Lu, Y., Tai, Y., She, G., Sun, J., Cao, H., Tong, W., Gao, Q., Li, Y., Deng, W., Jiang, X., Wang, W., Chen, Q., Zhang, S., Li, H., Wu, J., Wang, P., Li, P., Shi, C., Zheng, F., Jian, J., Huang, B., Shan, D., Shi, M., Fang, C., Yue, Y., Li, F., Li, D., Wei, S., Han, B., Jiang, C., Yin, Y., Xia, T., Zhang, Z., Bennetzen, J.L., Zhao, S., Wan, X., 2018. Draft genome sequence of *Camellia sinensis* var. *sinensis* provides insights into the evolution of the tea genome and tea quality. Proc. Natl. Acad. Sci. U. S. A.115, E4151–E4158. https://doi.org/10.1073/pnas.1719622115

70. Wei, K., Wang, L., Zhang, Y., Ruan, L., Li, H., Wu, L., Xu, L., Zhang, C., Zhou, X., Cheng, H., Edwards, R., 2019. A coupled role for *CsMYB75* and *CsGSTF1* in anthocyanin hyperaccumulation in purple tea. Plant J. 97, 825–840. https://doi.org/10.1111/tpj.14161

71. Wilkins, O., Nahal, H., Foong, J., Provart, N.J., Campbell, M.M., 2009. Expansion and diversification of the *Populus* R2R3-MYB family of transcription factors. Plant Physiol. 149, 981–993. https://doi.org/10.1104/pp.108.132795

72. Yamashita, H., Uchida, T., Tanaka, Y., Katai, H., Nagano, A.J., Morita, A., Ikka, T., 2020. Genomic predictions and genome-wide association studies based on RAD-seq of quality-related metabolites for the genomics-assisted breeding of tea plants. Sci. Rep. 10, 17480. https://doi.org/10.1038/s41598-020-74623-7

73. Yang, D., Liu, Y., Sun, M., Zhao, L., Wang, Y., Chen, X., Wei, C., Gao, L., Xia, T., 2012. Differential gene expression in tea (*Camellia sinensis* L.) calli with different morphologies and catechin contents. J. Plant Physiol. 169, 163–175. https://doi.org/https://doi.org/10.1016/j.jplph.2011.08.015

74. Yokozawa, T., Dong, E., Nakagawa, T., Kashiwagi, H., Nakagawa, H., Takeuchi, S., Chung, H.Y., 1998. In Vitro and in Vivo studies on the radical-scavenging activity of tea. J. Agric. Food Chem. 46, 2143–2150. https://doi.org/10.1021/jf970985c

75. Yu, M., Man, Y., Lei, R., Lu, X., Wang, Y., 2020. Metabolomics study of flavonoids and anthocyanin-related gene analysis in kiwifruit (*Actinidia chinensis*) and kiwiberry (*Actinidia arguta*). Plant Mol. Biol. Report. 38, 353–369. https://doi.org/10.1007/s11105-020-01200-7

76. Yu, S., Li, P., Zhao, X., Tan, M., Ahmad, M.Z., Xu, Y., Tadege, M., Zhao, J., 2021. *CsTCP*s regulate shoot tip development and catechin biosynthesis in tea plant (*Camellia sinensis*). Hortic. Res. 8, 104. https://doi.org/10.1038/s41438-021-00538-7

77. Yu, X., Xiao, J., Chen, S., Yu, Y., Ma, J., Lin, Y., Li, R., Lin, J., Fu, Z., Zhou, Q., Chao, Q., Chen, L., Yang, Z., Liu, R., 2020. Metabolite signatures of diverse *Camellia sinensis* tea populations. Nat. Commun. 11, 5586. https://doi.org/10.1038/s41467-020-19441-1

78. Zhang, P., Chopra, S., Peterson, T., 2000. A segmental gene duplication generated differentially expressed *myb*-homologous genes in Maize. Plant Cell 12, 2311. https://doi.org/10.2307/3871231

79. Zhang, Y., Butelli, E., Martin, C., 2014. Engineering anthocyanin biosynthesis in plants. Curr. Opin. Plant Biol. 19, 81–90. https://doi.org/10.1016/j.pbi.2014.05.011

80. Zhu, Z., Sun, B., Cai, W., Zhou, X., Mao, Y., Chen, Chengjie, Wei, J., Cao, B., Chen, Changming, Chen, G., Lei, J., 2019. Natural variations in the MYB transcription factor *MYB31* determine the evolution of extremely pungent peppers. New Phytol. https://doi.org/10.1111/nph.15853

