## Supplementary Table S1 to S4 for "Systematic analysis of the R2R3-MYB family of transcription factors in *Camellia sinensis*: evidence for species-specific catechin biosynthesis regulation"

**Supplementary Table S1**. Detailed subgroup situation of R2R3-MYB proteins in Figure 1

| Gene ID | Name | Subgroup | Function |
|---|---|---|---|
| AT1G34670 | AtMYB93 | Subgroup 10 & Subgroup 24 | Enable DNA-binding transcription factor activity, transcription cis-regulatory region binding |
| AT5G65230 | AtMYB53 | | |
| AT5G10280 | AtMYB92 | | |
| AT3G02940 | AtMYB107 | | |
| AT5G16770 | AtMYB9 | | |
| AT4G17785 | AtMYB39 | | |
| TEA007975 | CsMYB93d | | |
| TEA015814 | CsMYB93c | | |
| TEA012565 | CsMYB93a | | |
| TEA012539 | CsMYB93b | | |
| TEA003067 | CsMYB9d | | |
| TEA026389 | CsMYB9b | | |
| TEA001022 | CsMYB9c | | |
| TEA015155 | CsMYB9a | | |
| AT5G54230 | AtMYB49 | Subgroup 11 + AtMYB49 | Abiotic stress response |
| AT4G05100 | AtMYB74 | | |
| AT4G21440 | AtMYB102 | | |
| AT4G28110 | AtMYB41 | | |
| TEA001795 | CsMYB10b | | |
| TEA004932 | CsMYB102a | | |
| TEA032908 | CsMYB41 | | |
| TEA023311 | CsMYB102c | | |
| TEA026096 | CsMYB102d | | |
| AT1G18570 | AtMYB51 | Subgroup 12 | Glycosylate biosynthesis |
| AT1G74080 | AtMYB122 | | |
| AT5G60890 | AtMYB34 | | |
| AT5G07690 | AtMYB29 | | |
| AT5G07700 | AtMYB76 | | |
| AT5G61420 | AtMYB28 | | |
| AT1G18710 | AtMYB47 | AtMYB47 & AtMYB95 | Enable DNA-binding transcription factor activity |
| AT1G74430 | AtMYB95 | | |
| AT3G61250 | AtMYB17 | Subgroup 9b | Regulation of early inflorescence development and seed germination |
| TEA030893 | CsMYB17a | | |
| TEA016497 | CsMYB17b | | |

| | | | |
|---|---|---|---|
| AT5G15310<br>AT3G01140<br>TEA031473<br>TEA023584<br>TEA026496<br>TEA030475 | AtMYB16<br>AtMYB106<br>CsMYB106a<br>CsMYB16<br>CsMYB106b<br>CsMYB106c | Subgroup 9a | AtMYB16/MIXTA: cell fate / Conical epidermal cell outgrowth; AtMYB106/NOK: cell fate / Trichome branching |
| AT2G36890<br>AT5G23000<br>AT4G37780<br>AT5G57620<br>AT3G49690<br>AT5G65790<br>TEA021984<br>TEA012515<br>TEA018194<br>TEA025993<br>TEA029960<br>TEA015798<br>TEA011993<br>TEA021422<br>TEA009432<br>TEA014063 | AtMYB38<br>AtMYB37<br>AtMYB87<br>AtMYB36<br>AtMYB84<br>AtMYB68<br>CsMYB36e<br>CsMYB36f<br>CsMYB36d<br>CsMYB36a<br>CsMYB36b<br>CsMYB36c<br>CsMYB84a<br>CsMYB84b<br>CsMYB38<br>CsMYB36g | Subgroup14 | Axillary meristem regulation / Lateral organ formation (shoot branching, GA - mediated) |
| AT3G28470<br>AT5G56110<br>TEA028555<br>TEA028188<br>TEA023344 | AtMYB35<br>AtMYB103<br>CsMYB35a<br>CsMYB35b<br>CsMYB103 | Subgroup AtMYB35 & AtMYB103 | AtMYB35: anther wall tapetum development; AtMYB103: regulation of secondary cell wall biogenesis |
| AT5G14340<br>AT5G62320<br>AT1G66230<br>AT5G16600<br>AT4G22680<br>AT4G12350<br>TEA003248<br>TEA028963<br>TEA014701<br>TEA023817<br>TEA030065<br>TEA033570 | AtMYB40<br>AtMYB99<br>AtMYB20<br>AtMYB43<br>AtMYB85<br>AtMYB42<br>CsMYB40a<br>CsMYB40b<br>CsMYB85c<br>CsMYB20<br>CsMYB85b<br>CsMYB85a | SubgroupAtMYB20, AtMYB40, AtMYB42, AtMYB43, AtMYB85 & AtMYB99 | Regulation of lignin biosynthetic process, phenylpropanoid metabolic process and secondary cell wall biogenesis |
| AT5G26660 | AtMYB86 | Subgroup13 | Mucilage deposition and |

| | | | |
|---|---|---|---|
| AT1G09540 | AtMYB61 | | extrusion; |
| AT1G57560 | AtMYB50 | | Phenylpropanoid |
| AT4G01680 | AtMYB55 | | pathway / Lignin |
| TEA020034 | CsMYB86a | | biosynthesis; |
| TEA022787 | CsMYB86b | | Stomatal closure |
| TEA002399 | CsMYB61b | | |
| TEA033203 | CsMYB61a | | |
| AT3G13890 | AtMYB26 | SubgroupAtMYB26, AtMYB67 | Anther dehiscence, |
| AT3G12720 | AtMYB67 | & AtMYB103 | plant-type secondary |
| AT1G63910 | AtMYB103 | | cell wall biogenesis |
| TEA028875 | CsMYB26 | | |
| TEA021737 | CsMYB103b | | |
| TEA007243 | CsMYB103a | | |
| TEA015433 | CsMYB67 | | |
| AT5G12870 | AtMYB46 | AtMYB46 & AtMYB83 | Regulation of secondary |
| AT3G08500 | AtMYB83 | | cell wall biogenesis |
| TEA030156 | CsMYB46 | | |
| AT3G48920 | AtMYB45 | Subgroup 16 | Hypostyle elongation, |
| AT4G25560 | AtMYB18 | | Far red light - mediated |
| AT5G52260 | AtMYB19 | | (phyA signaling) |
| TEA017151 | CsMYB45 | | |
| TEA029135 | CsMYB19 | | |
| TEA027012 | CsMYB2 | TEA Preferential Subgroup A | |
| TEA033589 | CsMYB3 | | |
| TEA006889 | CsMYB5 | | |
| TEA033593 | CsMYB8 | | |
| TEA017225 | CsMYB10 | | |
| TEA022451 | CsMYB11 | | |
| AT1G08810 | AtMYB60 | Subgroup 1 | Abiotic and biotic stress |
| AT5G62470 | AtMYB96 | | response |
| AT3G28910 | AtMYB30 | | |
| AT3G47600 | AtMYB94 | | |
| AT1G74650 | AtMYB31 | | |
| TEA000749 | CsMYB94a | | |
| TEA023891 | CsMYB96 | | |
| TEA002289 | CsMYB60 | | |
| TEA014430 | CsMYB94b | | |
| AT1G16490 | AtMYB58 | Subgroup 3 & AtMYB10, | Phenylpropanoid |

| | | | |
|---|---|---|---|
| AT1G56160 | AtMYB72 | AtMYB72 | pathway / Lignin |
| AT1G79180 | AtMYB63 | | biosynthesis (fibers and |
| AT3G12820 | AtMYB10 | | vessels) |
| TEA023193 | CsMYB63 | | |
| AT1G06180 | AtMYB13 | Subgroup 2 | Abiotic stress response |
| AT2G31180 | AtMYB14 | | |
| AT3G23250 | AtMYB15 | | |
| TEA029615 | CsMYB15a | | |
| TEA019409 | CsMYB14a | | |
| TEA020748 | CsMYB15b | | |
| TEA028476 | CsMYB14b | | |
| TEA029605 | CsMYB15c | | |
| TEA029352 | CsMYB15d | | |
| AT4G38620 | AtMYB4 | Subgroup 4 & AtMYB6, | Abiotic stress responses |
| AT2G16720 | AtMYB7 | AtMYB8 | |
| AT1G22640 | AtMYB3 | | |
| AT1G35515 | AtMYB8 | | |
| AT4G09460 | AtMYB6 | | |
| AT4G34990 | AtMYB32 | | |
| TEA000547 | CsMYB4b | | |
| TEA003515 | CsMYB4a | | |
| TEA019219 | CsMYB7a | | |
| TEA008298 | CsMYB6 | | |
| TEA023057 | CsMYB7b | | |
| TEA025886 | CsMYB13 | TEA Preferential Subgroup B | |
| TEA033191 | CsMYB18 | | |
| TEA023874 | CsMYB22 | | |
| TEA032503 | CsMYB23 | | |
| AT5G49330 | AtMYB111 | Subgroup 7 | Phenylpropanoid |
| AT2G47460 | AtMYB12 | | pathway / Anthocyanin |
| AT3G62610 | AtMYB11 | | biosynthesis |
| TEA009412 | CsMYB12 | | |
| AT3G13540 | AtMYB5 | Subgroup AtMYB5 | Redundant with |
| TEA014311 | CsMYB37 | | AtMYB123 in regulating |
| TEA027333 | CsMYB39 | | tannin biosynthesis |
| TEA012130 | CsMYB42 | | |
| TEA012145 | CsMYB43 | | |
| AT3G29020 | AtMYB110 | Subgroup 21 + AtMYB89 | Axillary meristem |

| | | | |
|---|---|---|---|
| AT5G39700 | AtMYB89 | | regulation; |
| AT1G17950 | AtMYB52 | | Cell wall thickening |
| AT1G73410 | AtMYB54 | | |
| AT4G33450 | AtMYB69 | | |
| AT5G17800 | AtMYB56 | | |
| AT1G26780 | AtMYB117 | | |
| AT1G69560 | AtMYB105 | | |
| TEA021056 | CsMYB117a | | |
| TEA016361 | CsMYB105 | | |
| TEA027440 | CsMYB56 | | |
| TEA002233 | CsMYB117b | | |
| AT3G55730 | AtMYB109 | Subgroup 23 | Abscisic acid response |
| AT2G39880 | AtMYB25 | | |
| AT3G09230 | AtMYB1 | | |
| TEA013753 | CsMYB109 | | |
| TEA018401 | CsMYB1a | | |
| AT2G23290 | AtMYB70 | Subgroup22 | Abiotic stress response |
| AT4G37260 | AtMYB73 | | |
| AT3G50060 | AtMYB77 | | |
| AT5G67300 | AtMYB44 | | |
| TEA013585 | CsMYB73b | | |
| TEA014193 | CsMYB73a | | |
| TEA000452 | CsMYB44 | | |
| AT2G37630 | AtMYB91 | SubgroupAtMYB91 | Regulation of innate immune response |
| AT1G14350 | AtMYB124 | SubgroupAtMYB88&AtMYB124 | Guard cell differentiation |
| AT2G02820 | AtMYB88 | | |
| AT3G60460 | AtMYB125 | Subgroup AtMYB125 | Stamen development |
| TEA023292 | CsMYB125 | | |
| AT2G25230 | AtMYB100 | Subgroup 25 + AtMYB98 | Embryogenesis / Seed maturation |
| AT5G40430 | AtMYB22 | | |
| AT3G27785 | AtMYB118 | | |
| AT5G40360 | AtMYB115 | | |
| AT4G18770 | AtMYB98 | | |
| AT5G11050 | AtMYB64 | | |
| AT5G58850 | AtMYB119 | | |
| TEA013946 | CsMYB98 | | |

| | | | |
|---|---|---|---|
| AT2G26950<br>AT2G26960<br>AT3G11440<br>AT5G06100<br>AT4G26930<br>AT5G55020<br>AT2G32460<br>TEA021145<br>TEA025308<br>TEA028392 | AtMYB104<br>AtMYB81<br>AtMYB65<br>AtMYB33<br>AtMYB97<br>AtMYB120<br>AtMYB101<br>CsMYB33<br>CsMYB65<br>CsMYB101 | Subgroup 18 | Cell differentiation; Stamen development / Anther development; Abiotic stress response / ABA sensitivity |
| AT1G66370<br>AT1G66390<br>AT1G56650<br>AT1G66380<br>TEA018834 | AtMYB113<br>AtMYB90<br>AtMYB75<br>AtMYB114<br>CsAN1 | Subgroup 6 | Phenylpropanoid pathway / Anthocyanin biosynthesis |
| TEA000825<br>TEA004616<br>TEA001090<br>TEA029017 | CsMYB25<br>CsMYB28<br>CsMYB29<br>CsMYB30 | TEA Preferential Subgroup C | |
| AT5G52600<br>TEA023420 | AtMYB82<br>CsMYB82 | SubgroupAtMYB82 | Trichome differentiation |
| AT5G40330<br>AT3G27920<br>AT5G14750<br>TEA005813 | AtMYB23<br>AtMYB0<br>AtMYB66<br>CsMYB66 | Subgroup15 | Cell fate / Trichome initiation |
| AT3G53200<br>AT3G46130<br>AT5G59780<br>TEA031915<br>TEA005413<br>TEA012256 | AtMYB27<br>AtMYB48<br>AtMYB59<br>CsMYB27<br>CsMYB48b<br>CsMYB48a | SubgroupAtMYB27, AtMYB48 & AtMYB59 | Regulation of potassium ion transport |
| AT3G24310<br>AT4G13480<br>AT3G30210<br>TEA005395<br>TEA016845<br>TEA028103 | AtMYB71<br>AtMYB79<br>AtMYB121<br>CsMYB71b<br>CsMYB71a<br>CsMYB121 | Subgroup AtMYB71, AtMYB79 & AtMYB121 | Enable DNA-binding transcription factor activity, transcription cis-regulatory region binding |

| | | | |
|---|---|---|---|
| AT5G35550 | AtMYB123 | Subgroup 5 | Phenylpropanoid pathway / Proanthocyanin dins biosynthesis |
| TEA004608 | CsMYB31 | | |
| TEA002308 | CsMYB32 | | |
| TEA031375 | CsMYB34 | | |
| AT3G01530 | AtMYB57 | Subgroup 19 + AtMYB57 | Stamen development / Filament length, GA‐ and JA‐mediated |
| AT3G27810 | AtMYB21 | | |
| AT5G40350 | AtMYB24 | | |
| TEA030389 | CsMYB24 | | |
| TEA024596 | CsMYB21 | | |
| AT1G48000 | AtMYB112 | Subgroup20 | Abiotic and biotic stress response |
| AT5G49620 | AtMYB78 | | |
| AT3G06490 | AtMYB108 | | |
| AT2G47190 | AtMYB2 | | |
| AT1G25340 | AtMYB116 | | |
| AT1G68320 | AtMYB62 | | |
| TEA012423 | CsMYB108 | | |
| TEA024384 | CsMYB78 | | |
| TEA013109 | CsMYB116 | | |
| TEA027578 | CsMYB62a | | |
| TEA002040 | CsMYB1 | | |
| TEA002210 | CsMYB62b | | |

**Supplementary Table S2**. List of qPCR primers used in this study

| Gene ID | Gene name | Primers |
|---|---|---|
| TEA027012 | CsMYB2 | qCsMYB2-F: TGGCGTGAACTTCCCAATGA |
| | | qCsMYB2-R: TTCCCCGTTTCACGTTTGGA |
| TEA033589 | CsMYB3 | qCsMYB3-F: TGGTCATTGGAACTGGCGTG |
| | | qCsMYB3-R: GCTTTGGTGTACTTTCCCCG |
| TEA006889 | CsMYB5 | qCsMYB5-F: ACCTGCGTCCCAATGTGAAA |
| | | qCsMYB5-R: ATCGCTTCTTGAGGTGGGTG |
| TEA033593 | CsMYB8 | qCsMYB8-F: GCGCAAAAACCACGCTCTAA |
| | | qCsMYB8-R: GTGGGGGTGAGAATTGGGAG |
| TEA017225 | CsMYB10 | qCsMYB10-F: CACACCCACCTGAAAAAGCG |
| | | qCsMYB10-R: CCCAATAGCAGGGCCAGTAG |
| TEA022451 | CsMYB11 | qCsMYB11-F: AAATGGTCTGCAATCGCTGC |
| | | qCsMYB11-R: ATGCTTGTGATTGGGTCCGT |
| TEA025886 | CsMYB13 | qCsMYB13-F: CCAGGGCGAACAGACAATGA |
| | | qCsMYB13-R: CATCCGACCCAGACGAAGTG |
| TEA033191 | CsMYB18 | qCsMYB18-F: CGGTGGTCTCTGATAGCTGG |
| | | qCsMYB18-R: AACCCACATGACACAAGGGG |
| TEA023874 | CsMYB22 | qCsMYB22-F: GGCCTACTTCGTTGCGGTAA |
| | | qCsMYB22-R: GTTCGTCCTGGCAATCTCCC |
| TEA032503 | CsMYB23 | qCsMYB23-F: GCTGCAGGCTTACTTCGTTG |
| | | qCsMYB23-R: ATCCGTTCGTCCTGGCAATC |
| TEA000825 | CsMYB25 | qCsMYB25-F: TGGCATCCAAGAGAACTGAGTC |
| | | qCsMYB25-R: GCACCATGGCTTTCGATGAC |
| TEA004616 | CsMYB28 | qCsMYB28-F: TGGCGGCTCCTAAGAACAAC |
| | | qCsMYB28-R: GATTCCCAGCGTTTTGAGCC |
| TEA001090 | CsMYB29 | qCsMYB29-F: ACCAATGGGTTGGAATGGGT |
| | | qCsMYB29-R: CCGGTGAAGGTGGAGTTCAA |
| TEA029017 | CsMYB30 | qCsMYB30-F: GGTCCTTGATTGCTGGGAGA |
| | | qCsMYB30-R: CGGCGAAGTTGTCGTCATTT |
| TEA004608 | CsMYB31 | qCsMYB31-F: CCGACCGCAGTCAAGAAGAA |
| | | qCsMYB31-R: TTCAAGCCTTGGTTGGGAGG |
| TEA002308 | CsMYB32 | qCsMYB32-F: AGTCTCTGCAATGGGTTGCT |
| | | qCsMYB32-R: TCCGACAAGCAAACATCCCT |
| TEA031375 | CsMYB34 | qCsMYB34-F: ACCAAGCGCAACCTCAACTA |
| | | qCsMYB34-R: GTACGGACTAGGCGTGATGG |
| TEA014311 | CsMYB37 | qCsMYB37-F: TCTTGGCAACCGATGGTCTC |
| | | qCsMYB37-R: TATTCGGGTCGGTTCCTTGG |
| TEA027333 | CsMYB39 | qCsMYB39-F: TTATCGCTGGAAGGCTACCG |
| | | qCsMYB39-R: GGGCATTGGGTTCTGTTCCT |
| TEA012130 | CsMYB42 | qCsMYB42-F: GCGGATGACAAACCGTCTTC |
| | | qCsMYB42-R: CGAGGTTTGCTCCCTCAGAA |
| TEA012145 | CsMYB43 | qCsMYB43-F: AGCCGGTCAGTAGTAGTGGT |

| | | |
|---|---|---|
| | | qCsMYB43-R: ATTGCAGCCACCAAGACCAT |
| TEA014056 | CsPAL | qPAL-F: GAAGGCCTTGCATTAGTGAATG |
| | | qPAL-R: TATGGGTCAAATGGTCCGTAAA |
| TEA026294 | CsF3'5'H | qF3'5'H-F: AAAGTAATTGGAAGAAACCGCC |
| | | qF3'5'H-R: TAGGAATACTGAGGGGAACTGA |
| TEA023790 | CsF3H | qF3H-F: GACCTACTTCTCATACCCGATC |
| | | qF3H-R: AGTCCATCAATTTCTCGCTGTA |
| TEA027582 | CsLAR | qLAR-F: GTGTTGGAATCTGTGTCCGCAG |
| | | qLAR-R: CTCCATGAATGCTTGATCCTTG |
| TEA010322 | CsANS | qANS-F: CTCCATCGTGGACTCGTTAATA |
| | | qANS-R: AGAATGATCTTCTCCTTGGGTG |
| TEA022960 | CsANR | qANR-F: AACCAGCAATTCAAGGAGTAGT |
| | | qANR-R: TCCCATTGAGCTTATTGATCGA |
| TEA006643 | CsFLS | qFLS-F2: ATGGAGGTAGAGAGAGTGCAAGCC |
| | | qFLS-R2: GTGGTTGAGAGAGGGAGATCACGG |
| TEA032730 | CsDFR | qDFR-F: ATGAAAGACTCTGTTGCTTCTG |
| | | qDFR-R: TGCTTCACCTTCTTTAAATTCG |
| TEA014864 | CsC4H | qC4H-F: GGCATAGCAGAACTTGTAAACC |
| | | qC4H-R: TTTGACTACAGCTTGGAGGTAG |
| TEA025906 | Cs4CL | q4CL-F: GATGTGATCATGTGTGTGCTAC |
| | | q4CL-R: CGCAATCGTCACCTTATACTTG |
| TEA009664 | CsSCPL1A7 | qSCPL1A7-F2: CGCGCCAATATACTATATGTAGAT |
| | | qSCPL1A7-R2: ATAAGAATCACCACCAATGTATAG |

**Supplementary Table S3**. The data acquired by HPLC system

| **Catechins** | **AB** | **FL** | **SL** | **ML** | **OL** | **S** | **R** |
|---|---|---|---|---|---|---|---|
| EGC | 20.67 | 20.85 | 26.05 | 26.88 | 8.49 | 1.95 | 0.00 |
| C | 1.65 | 1.67 | 1.24 | 0.86 | 0.25 | 5.54 | 0.00 |
| EC | 4.76 | 5.13 | 4.78 | 4.56 | 3.14 | 0.80 | 1.80 |
| EGCG | 124.12 | 125.44 | 119.28 | 104.39 | 76.34 | 44.30 | 0.52 |
| ECG | 23.99 | 23.15 | 19.14 | 12.96 | 10.39 | 10.26 | 0.29 |

Note: The contents of EGC, C, EC, EGCG and ECG (mg·g-1 dry weight) were detected in 7 different tissues of tea cultivar Lingtoudancong using HPLC system. The studied 7 tissues were apical buds (AB), first leaves (FL), second leaves (SL), mature leaves (ML), old leaves (OL), young stems (S), and tender roots (R).

**Supplementary Table S4.** Expression profiles of CsR2R3-MYB genes and structural genes in different tea plant tissues

| Gene names | AB | FL | SL | ML | OL | S | R |
|---|---|---|---|---|---|---|---|
| CsPAL | 0.05308 | 0.09189 | 0.12247 | 0.31522 | 0.18081 | 0.11479 | 0.28362 |
| CsF3'5'H | 0.01078 | 0.01104 | 0.00932 | 0.01110 | 0.00073 | 0.00651 | 0.00148 |
| CsF3H | 0.06118 | 0.04176 | 0.04553 | 0.04442 | 0.00705 | 0.02132 | 0.01177 |
| CsLAR | 0.98791 | 0.85824 | 0.94610 | 0.88065 | 0.00082 | 0.17009 | 0.13420 |
| CsANS | 0.01448 | 0.00952 | 0.00484 | 0.00298 | 0.00024 | 0.03728 | 0.00521 |
| CsANR | 0.60796 | 0.49734 | 0.66477 | 0.43280 | 0.14094 | 0.45716 | 0.18583 |
| CsFLS | 0.72917 | 1.26354 | 1.07825 | 0.81241 | 0.10463 | 0.48638 | 0.34021 |
| CsDFR | 0.03382 | 0.02310 | 0.01539 | 0.00639 | 0.00152 | 0.01381 | 0.00354 |
| CsC4H | 0.08643 | 0.09382 | 0.13892 | 0.19560 | 0.11005 | 0.12122 | 0.73786 |
| Cs4CL | 0.07179 | 0.04127 | 0.04432 | 0.07572 | 0.02568 | 0.03366 | 0.12495 |
| CsSCPL1A7 | 0.31013 | 0.14847 | 0.05521 | 0.01132 | 0.00407 | 0.00992 | 0.00689 |
| CsMYB2 | 0.00015 | 0.00013 | 0.00016 | 0.00065 | 0.00948 | 0.00010 | 0.00000 |
| CsMYB3 | 0.01152 | 0.01253 | 0.01321 | 0.00670 | 0.03316 | 0.00832 | 0.01343 |
| CsMYB5 | 0.00831 | 0.00202 | 0.00320 | 0.00813 | 0.02877 | 0.00056 | 0.00257 |
| CsMYB8 | 0.43701 | 0.22453 | 0.39124 | 0.28611 | 0.06249 | 0.19460 | 0.07373 |
| CsMYB10 | 0.00003 | 0.00012 | 0.00022 | 0.00009 | 0.01620 | 0.00002 | 0.01716 |
| CsMYB11 | 0.00116 | 0.00027 | 0.00227 | 0.00288 | 0.01860 | 0.00044 | 0.01434 |
| CsMYB13 | 0.04363 | 0.00987 | 0.02210 | 0.13524 | 0.16418 | 0.00784 | 0.03903 |
| CsMYB18 | 0.00022 | 0.00009 | 0.00099 | 0.00050 | 0.00114 | 0.00012 | 0.00004 |
| CsMYB22 | 0.12403 | 0.06590 | 0.10940 | 0.16820 | 0.34881 | 0.02667 | 0.05946 |
| CsMYB23 | 0.08193 | 0.04078 | 0.28359 | 0.18949 | 0.08244 | 0.01356 | 0.02862 |
| CsMYB25 | 0.01384 | 0.00826 | 0.01382 | 0.03597 | 0.02335 | 0.00163 | 0.00242 |
| CsMYB28 | 0.01545 | 0.00294 | 0.02095 | 0.01039 | 0.00604 | 0.00142 | 0.00117 |
| CsMYB29 | 0.00098 | 0.00093 | 0.00087 | 0.00006 | 0.00075 | 0.00007 | 0.00054 |
| CsMYB30 | 0.22921 | 0.06104 | 0.12926 | 0.04335 | 0.00226 | 0.00599 | 0.00811 |
| CsMYB31 | 0.01257 | 0.00461 | 0.00997 | 0.02099 | 0.05942 | 0.00084 | 0.01403 |
| CsMYB32 | 0.00864 | 0.00455 | 0.00375 | 0.00987 | 0.00309 | 0.00041 | 0.00158 |
| CsMYB34 | 0.10623 | 0.04879 | 0.05211 | 0.04053 | 0.00404 | 0.02227 | 0.00752 |
| CsMYB37 | 0.41759 | 0.38520 | 0.21531 | 0.29008 | 0.02953 | 0.15456 | 0.03107 |
| CsMYB39 | 0.00672 | 0.00706 | 0.00452 | 0.00017 | 0.00746 | 0.00407 | 0.00085 |
| CsMYB42 | 0.05665 | 0.02804 | 0.02312 | 0.02625 | 0.10370 | 0.02057 | 0.00866 |
| CsMYB43 | 0.02818 | 0.02981 | 0.02716 | 0.03482 | 0.08283 | 0.02661 | 0.00185 |

Note: The studied 7 tissues were AB, apical buds; FL, first leaves; SL, second leaves; ML, mature leaves; OL, old leaves; S, young stems and R, tender roots.
